# Pre-Implantation Alcohol Exposure Induces Lasting Sex-Specific DNA Methylation Programming Errors in the Developing Forebrain

**DOI:** 10.1101/2020.12.07.415208

**Authors:** LM Legault, K Doiron, M Breton-Larrivée, A Langford-Avelar, A Lemieux, M Caron, LA Jerome-Majewska, D Sinnett, S McGraw

## Abstract

**Background:** Prenatal alcohol exposure is recognized for altering DNA methylation profiles of brain cells during development, and to be part of the molecular basis underpinning Fetal Alcohol Spectrum Disorder (FASD) etiology. However, we have negligible information on the effects of alcohol exposure during pre-implantation, the early embryonic window marked with dynamic DNA methylation reprogramming, and on how this may rewire the brain developmental program.

**Results:** Using a pre-clinical *in vivo* mouse model, we show that a binge-like alcohol exposure during pre-implantation at the 8-cell stage leads to surge in morphological brain defects and adverse developmental outcomes during fetal life. Genome-wide DNA methylation analyses of fetal forebrains uncovered sex-specific alterations, including partial loss of DNA methylation maintenance at imprinting control regions, and abnormal *de novo* DNA methylation profiles in various biological pathways (e.g., neural/brain development).

**Conclusion:** These findings support that alcohol-induced DNA methylation programming deviations during pre-implantation could contribute to the manifestation of neurodevelopmental phenotypes associated with FASD.

## INTRODUCTION

Fetal Alcohol Spectrum Disorders (FASD) encompasses the range of lifelong cognitive and physical disabilities observed in children born to mothers who consumed alcohol during pregnancy (1–4). Each year, 600 000 to 1 million children are born with FASD worldwide (5, 6). The most severe and physically visible form of the condition is known as Fetal Alcohol Syndrome (FAS), in ~10% of FASD cases, and associated with the full presentation of dysmorphic features including craniofacial malformations, growth deficits and structural brain pathologies. Depending on the amount, pattern and developmental period of prenatal alcohol exposure, children may not present with dysmorphic features, but still suffer from mild to severe FASD-related neurological disabilities, such as learning deficits and intellectual delays (1, 2). With the marked increase in rates of alcohol use and binge drinking behavior (7) among 18-34 year old women (8–14), added to the high number of unintended pregnancies worldwide (~40%; 85 million/year (15)), many women may inadvertently subject their developing embryos to acute levels of alcohol in first weeks of pregnancy. Although most studies state that alcohol consumption at all stages of pregnancy can cause FASD, pre-implantation is arguably the stage that is most prone to unintentional prenatal alcohol exposure since the human chorionic gonadotrophin (hCG) hormone, the main biomarker for pregnancy, is not yet detectable as it is only produced following implantation. Still, there is ample misinformation in the literature regarding the effects of alcohol exposure, and many other teratogen exposures, on pre-implantation embryos, and how this leads to an “*all-or-nothing*” developmental outcome.

An increasing body of evidence indicates that alcohol exposure during fetal brain development triggers lasting epigenetic alterations, including DNA methylation, in offspring long after the initial insult, supporting the role of epigenetics in FASD phenotypes (16–19). However, we remain unaware of how ethanol affects the early developmental window marked with dynamic changes in DNA methylation, and how interfering with this fundamental process may program future FASD-related neurological disabilities. During pre-implantation development, the period between oocyte fertilization and embryo implantation in the uterus, the epigenome undergoes a broad reprogramming that initiates the developmental program (20–26). We and others have shown that this essential reprogramming wave removes most DNA methylation signatures across the genome, except specific sequences that include imprinting control regions (ICRs), to trigger the embryonic developmental program (20, 27–29). DNA methylation marks are then reacquired in a sex-, cell- and tissue-specific manner during the peri-implantation period, and marks continue to be modulated during lineage specification (30–33). Studies show that pre-implantation embryos can have sex-specific epigenetic responses to similar environmental challenges, leading to long-term sexual dimorphism in developmental programming trajectories (34–37).

One of the first indications of the direct link between ethanol exposure and aberrations in DNA methylation came from a mouse study showing that mid-gestation exposure at E9-E11 reduced global DNA methylation levels in E11 fetuses (17). This evidence gave rise to different FASD models using various levels of short or prolonged alcohol exposures at different stages of gestation. Ethanol exposure can either induce DNA methyltransferases (DNMTs) activity, through reactive oxygen species-dependent mechanisms (38, 39), or inhibit DNMTs activity, via direct action on DNMTs or on one-carbon metabolism that provides methyl groups (17, 40), which supports why both gain and loss of DNA methylation marks can be observed in FASD models. High levels of ethanol exposure on two consecutive days (E1.5, E2.5) altered DNA methylation of imprinted gene *H19*, a negative regulator of growth and proliferation, in the placenta at E10.5, yet this region showed no alteration in the embryo (41). Nonetheless, data remain very scarce on how alcohol exposure during the early stages of embryo development directly affects epigenetic reprogramming and permanently alters genome-wide DNA methylation.

In this study, we used a pre-clinical mouse model of prenatal alcohol exposure to specifically target pre-implantation embryos that are undergoing their epigenetic reprogramming wave. We show that this exposure leads to a surge in morphological brain defects during fetal life, and that exposed embryos with no visible abnormalities or developmental delays present lasting DNA methylation alterations in forebrain tissues, including sex-specific disparities in DNA methylation dysregulation.

## RESULTS

### Modeling early pre-implantation alcohol exposure increases phenotypic alterations in developing embryos

To define the developmental and epigenetic (i.e., DNA methylation) impact of a binge alcohol exposure episode on early embryos undergoing the epigenetic reprogramming wave, we first established a pre-clinical mouse model to specifically expose pre-implantation embryos (E2.5) to short, but elevated alcohol levels. To avoid possible confounding effects of gavage-associated stress, we used a well-recognized two-injection paradigm (42–45). Pregnant mice (C57BL/6) were subcutaneously injected at E2.5 (8-cell embryos) with two doses of 2.5g/kg ethanol (**EtOH-exposed**), or 0.15M saline (**control**), at 2h intervals (**Fig. 1A**). Pregnant females reached a peak blood alcohol concentration (**BAC**) of 284.27mg/dL (3h) with an average of 158.31mg/dL over a 4h window (**Fig. 1B**). In contrast with other chronic prenatal alcohol exposure models (46–49), this short but acute level of ethanol exposure on pre-implantation embryos did not affect average mid-gestational (E10.5) litter size (Ctrl; n=8.13 ± 2.58 vs EtOH-exposed; n=7.91 ± 2.86), or sex distribution (Ctrl and EtOH-exposed; 49% female vs 51% males). Similarly, early pre-implantation embryos subjected to binge-like alcohol levels did not show differences in mean morphological measurements at E10.5, however, we observed a significant increase in embryo-to-embryo variability for crown to rump length (p<0.0001), head height (p<0.01), occipital to nose diameter (p<0.001) and brain sagittal length (p<0.05) for EtOH-exposed embryos compared to controls. The greater intra-subject variability observed suggests that binge alcohol exposure levels during pre-implantation can alter the normal developmental programming of early embryos, thus causing abnormal morphological outcomes.

**Figure 1.**
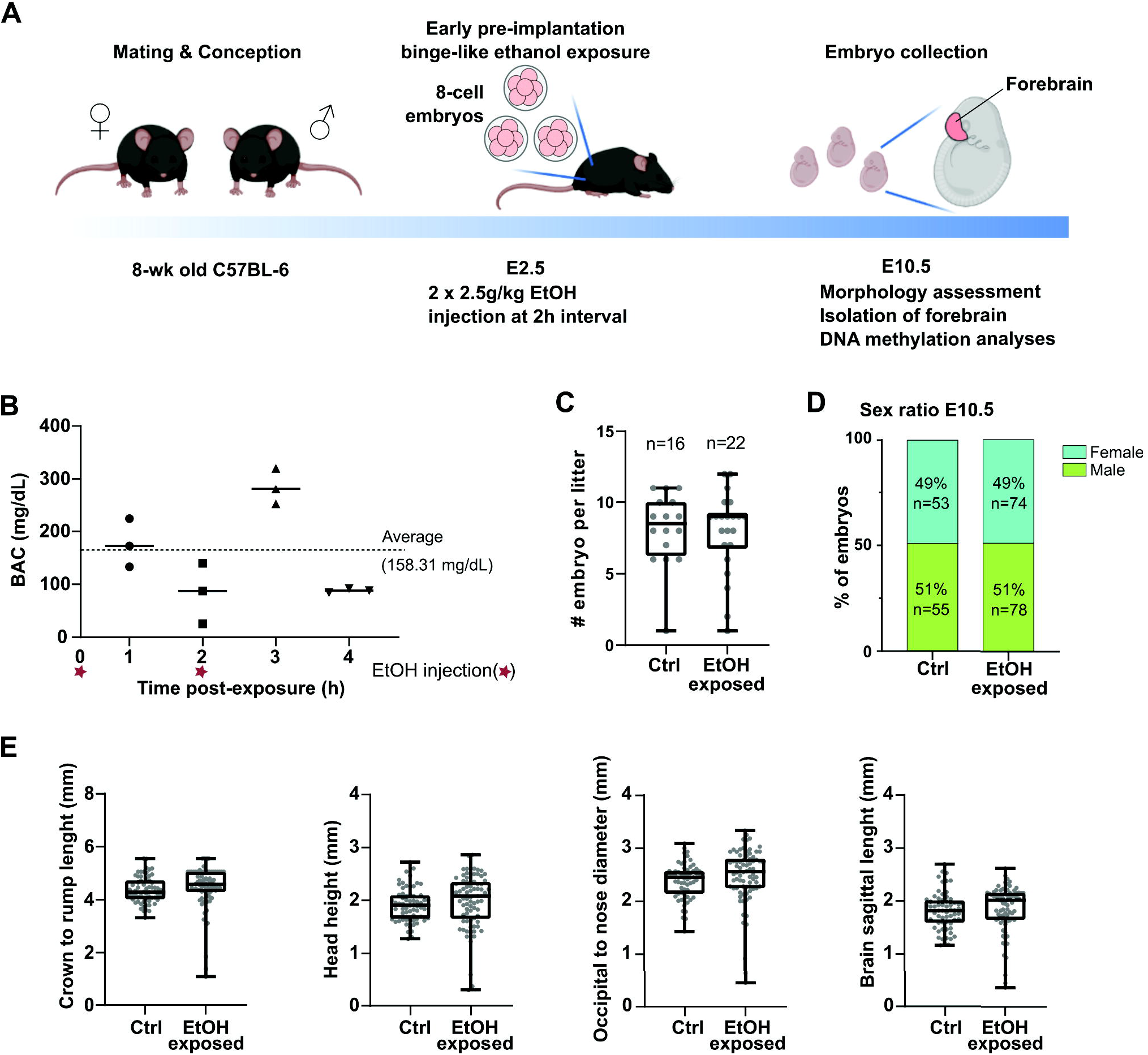
Early pre-implantation alcohol exposure affects mouse embryonic development at mid-gestation. **A)** Schematic description of binge alcohol exposure model during pre-implantation embryo development, and analyses performed at mid-gestation. Pregnant females were exposed to EtOH (2x 2.5g/kg EtOH) or Saline (Ctrl) (equivalent volume to ethanol) by subcutaneous injections (2h interval) to precisely target E2.5 stage embryos (~8-cell stage). E10.5 embryos were collected for morphological assessment; forebrain was isolated for genome-wide DNA methylation analyses. **B)** Quantification of blood alcohol concentration in pregnant females following EtOH exposure at E2.5. Red stars indicate the two EtOH injection time points. Peak level (284mg/dL) was observed at 3h post-exposure, with an average of 158.31 mg/dL over 4hrs. **C)** Number of E10.5 embryos per litter in Ctrl (n=16 litters; average 8.13 embryos/litter) and EtOH-exposed (n=22 litters; average 7.91 embryos/litter). **D)** Male and female embryo sex ratios of litters presented in panel C), with number of embryos shown in bar graph. **E)** Embryonic (E10.5) measurements: crown-rump length, head height, occipital-nose length and brain sagittal length. Control embryos: (n=63, 8 litters), ethanol-exposed embryos (n=76, 11 litters). No significant difference of the means; t-test with Welch’s correction, but higher variance in EtOH-exposed; F-test.

To define how a binge alcohol exposure episode during early pre-implantation can affect fetal development, we next investigated the morphological outcome of E10.5 embryos (control embryos: n=108 from 16 litters, EtOH-exposed: n=152 from 22 litters). As observed in **Fig. 2A**, early pre-implantation EtOH exposure leads to a significant increase in morphological defects or delayed development of mid-gestation embryos (19% vs 2%, p<0.0001). Types of defects observed in ethanol-exposed embryos included brain anomalies (e.g., forebrain or midbrain malformations) (10%), growth restriction or delayed development (5%), heart defects (2%), and other abnormal features (2%) (**Fig. 2B, Fig. 2C**). There was no sex-specific phenotypic divergence between EtOH-exposed male and female embryos for morphological defects (18% vs 20%, **Fig. S1**) or defect categories (**Fig. S1**). Compared to controls, we observed that binge alcohol exposure during pre-implantation leads to a larger proportion of E10.5 embryos with phenotypic alterations (29/152 vs 2/108, p<0.0001), and that embryos with phenotypic alterations are distributed across most ethanol-exposed litters (16/22 vs 2/16, p<0.001) (**Fig. 2D**) and not restricted to a small number of litters. Taken together, these results show that a binge alcohol exposure episode on pre-implantation embryos undergoing the epigenetic reprogramming wave does not interfere with normal processes of implantation but leads to heterogeneity in morphological presentation during fetal life that mirrors the spectrum of clinical features associated to FASD.

**Figure 2.**
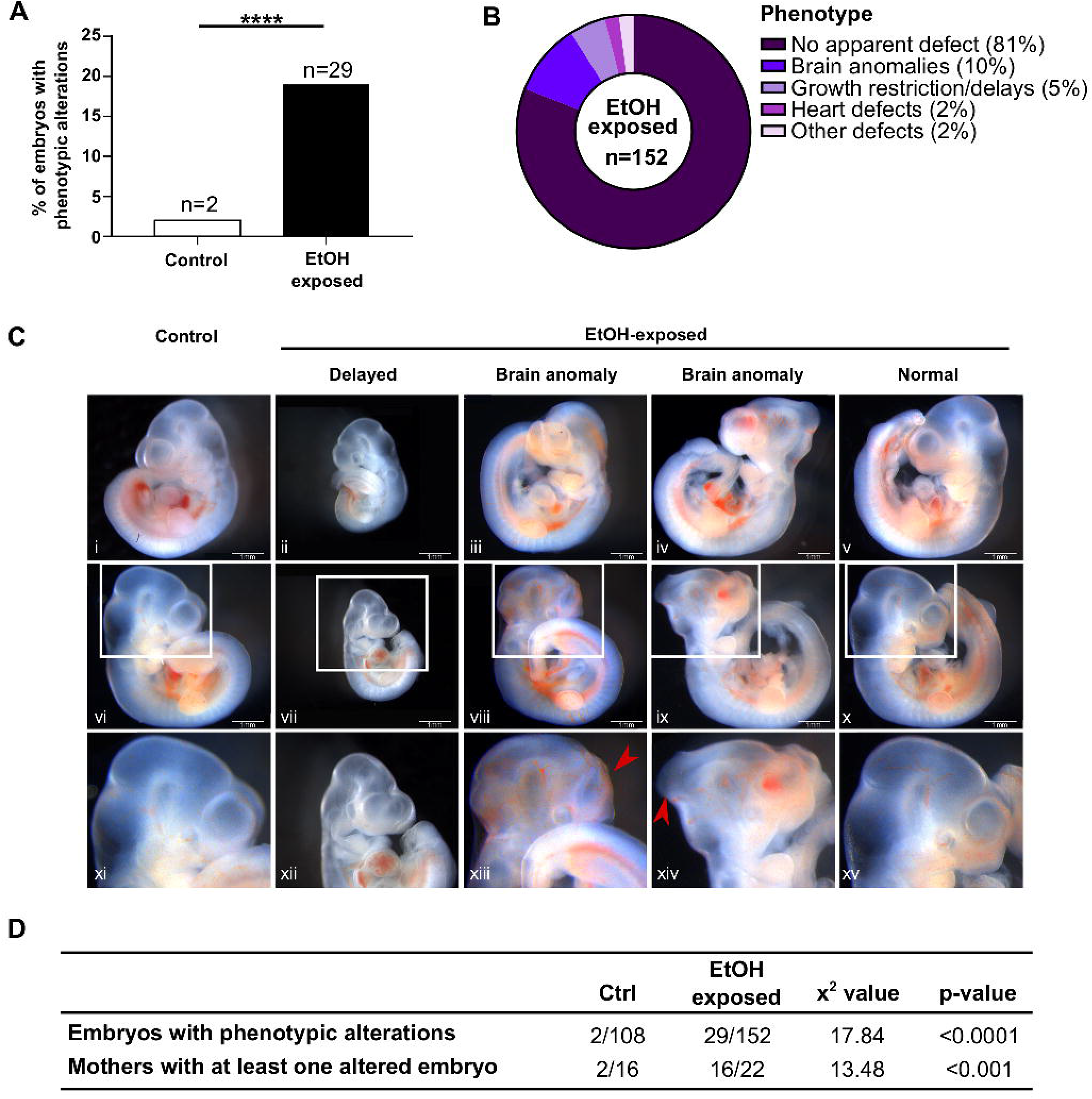
Increased phenotypic alterations in developing embryos following early pre-implantation alcohol exposure. **A)** Percentage of E10.5 embryos with phenotypic alterations (Ctrl: 2%; n=2/108, EtOH-exposed: 19%; n=29/152, ****p<0.0001; chi-square test). **B**) Classification and proportion of phenotypic alterations observed in EtOH-exposed embryos. **C)** Examples of primary phenotypic alterations observed in EtOH-exposed embryos. Views from both side of the embryo and zoom on the head for control (i, vi, xi) and EtOH-exposed embryos with severe developmental delays (ii, vii, xii), brain malformations (iii, viii, xiii, iv, ix, xiv) and no apparent defects (v, x, xv). **D)** Number of embryos with alterations and number of litters with at least one affected embryo. ***p<0.001; chi-square test.

### Pre-implantation alcohol exposure causes alterations in forebrain DNA methylation profiles

To assess whether alcohol exposure during the embryonic epigenetic reprogramming wave dysregulates the normal programming of brain DNA methylation patterns during fetal development, we established genome-wide DNA methylation profiles using rRRBS on E10.5 mouse forebrains. To mirror the 90% of children with FASD that show no dysmorphic features but still suffer from mild to severe neurological disabilities (e.g., learning deficits, intellectual delays), we randomly selected 6 controls (3 males; 3 females) and 16 ethanol-exposed embryos (8 males; 8 females) of similar size (i.e., embryo size, head height, occipital to nose diameter, brain sagittal length) with no visible morphological defects (Fig. **S2**). Histological analysis revealed that ethanol-exposed embryos with no visible morphological defects or developmental delays had a general layout and distribution of brain cells that were comparable to controls (**Fig. S3A-B**), whereas ethanol-exposed embryo with delayed development were distinctly different (**Fig. S3C**). Furthermore, using markers for proliferation (Ki67 antigen) and apoptosis (cleaved Caspase-3), we confirmed that the early ethanol exposure did not promote an imbalance in cell proliferative response (**Fig. S4**) or cell death (**Fig. S5**) across brain regions of embryos with no apparent morphological defects. By removing embryos with abnormalities, developmental delays, or gross brain structure aberration, we further reduced potential DNA methylation variability that could be due to divergent forebrain cell type proportions between samples.

The first set of analyses was designed to define whether early pre-implantation alcohol exposure caused lasting DNA methylation alterations in developing E10.5 embryonic forebrains despite the absence of phenotypic presentation. To do so, we compared the average DNA methylation levels in 100 bp non-overlapping genomic windows (*tiles*; see methods section) between controls and ethanol-exposed samples. After removal of sex chromosomes, we identified 114 911 unique sequenced tiles containing 794 803 common CpGs across samples (min. 5 samples/condition, ≥10x sequencing depth). When tiles were classified according to their DNA methylation levels, we observed significant changes in the distribution of categories, including tiles ranging between 90-100% methylation for which we observed a decrease in ethanol-exposed forebrains compared to controls (26% vs 24%, p<0.0001 (**Fig. 3A**). Clustering of individual E10.5 forebrain samples by DNA methylation levels for the top 1% most variable tiles (n=1200) revealed three main subgroups; a first essentially composed of control samples (6/9; right in heatmap), a second mainly composed of ethanol-exposed female samples with similarities to controls patterns (5/6; middle in heatmap), and a third mostly composed of male ethanol-exposed samples with highly divergent patterns (6/7; left in heatmap) (**Fig. 3B**). We next identified regions of the genome that showed altered DNA methylation levels (±≥ 10% mean differences between tiles; see methods) as a result of the early pre-implantation alcohol exposure. Using such criteria, we identified 1509 differentially methylations regions (**DMRs**) with significant DNA methylation level decrease (n=1440) or increase (n=69) in EtOH-exposed forebrains compared to control forebrains (**Fig. 3C**). Differences in methylation levels ranged from 10%-55%, with most DMRs showing 10-15% (n=1086; 72%) or 15-20% (n=300; 20%) methylation change between EtOH-exposed and control forebrains (**Fig. 3D**). Most DMR-associated tiles with decreased methylation levels in EtOH-exposed samples (n=1239) were highly methylated (≥40-50%) in control samples, whereas DMRs that gained methylation in EtOH-exposed samples had variable levels in controls (**Fig. 3E**). The DMRs mainly overlapped with intergenic (43%) and genic (53%: introns; 37%, exons; 12%, and promoters; 4%) regions. Gene ontology enrichment analyses associated genic DMRs with various processes, including cell-to-cell signaling pathway, regulation of GTPase activity, positive regulation of nervous system development, tissue and embryonic morphogenesis, as well as regulation of neurological systems (**Fig. 3F**). When we looked at the DNA methylation levels between control and EtOH-exposed forebrains for genes associated to these enriched pathways (e.g,. *Lrch1*, *Cflar*, *Celsr1*, *Apoa1*, *Lpin1*, *Epha7*, *Foxa1*, *Egf*, *Il6*, *Nkx6-2*), we observed increased inter-individual variability in methylation levels for EtOH-exposed forebrains with some samples or genomic regions being more affected than others (**Fig. 3G**).

**Figure 3.**
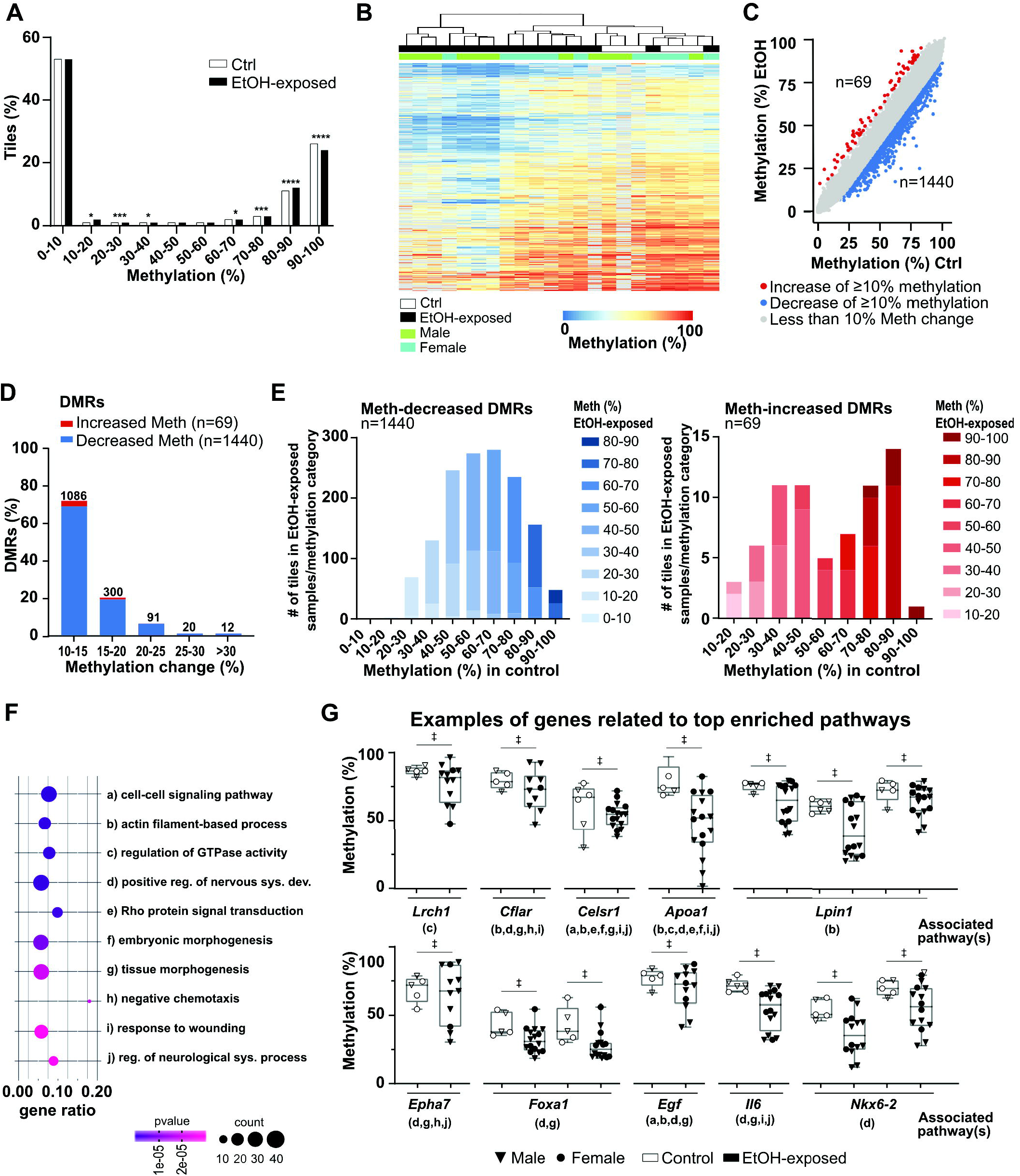
Early pre-implantation alcohol exposure triggers DNA methylation alterations in developing embryonic forebrain. Genome-wide CpG methylation analyses of E10.5 control (n=6) and ethanol-exposed (n=16) forebrains. **A)** Distribution of genomic tiles (100bp) (all-tiles; n=114 911) across ranges of CpG methylation levels in control and EtOH-exposed samples. ****p<0.0001, ***p<0.001, *p<0.05; z-test proportion test. **B)** Heatmap showing CpG methylation levels for the top1% most variable tiles (n=1 200) between control and EtOH-exposed forebrains. Gray lines in heatmap have no associated methylation values because of lack of sufficient sequencing coverage in sample. **C)** Scatterplot representing the differentially methylated regions (DMRs) between control and EtOH-exposed forebrains (see methods section for details). Red dots represent the tiles with a methylation increase of at least 10% in EtOH-exposed compared to control forebrains (n=69); blue dots represent the tiles with a methylation decrease of at least 10% in EtOH-exposed compared to control forebrains (n=1 440); grey dots represent the tiles with changes less than 10% in EtOH-exposed compared to control forebrains (n=113 402). **D)** Proportion of DMRs associated with the changes of CpG methylation levels between control and EtOH-exposed E10.5 forebrains. **E)** Comparison of CpG methylation levels of specific DMRs-associated tiles in EtOH-exposed versus control forebrains. Blue bar graph represents the comparison for decreased-methylation DMRs (n=1 440); red bar graph represents comparison for the increased-methylation DMRs (n=69). **F)** Functional enrichment analysis showing top 10 enriched pathways for decreased- and increased-methylation DMRs located in genic regions (n=710 unique gene DMRs), based on Metascape analysis for pathways and p-value. The size of the dot represents the number of DMR-associated genes in a pathway, and gene ratio represents the number of DMR-associated genes with regards to the number of genes in a pathway. **G)** Examples of CpG methylation levels of individual samples for DMR-associated genes related to the top enriched pathways. Letters under gene name relate to the pathways in **F).** □ represents significant differences in CpG methylation levels of DMRs (e.g., ±>10% methylation difference, q<0.01) between control and EtOH-exposed embryos (see methods section for details).

Overall, we showed that alcohol exposure during the embryonic epigenetic reprogramming wave triggers an array of DNA methylation alterations observed in forebrain of embryos that presented no visible abnormalities or developmental delays at E10.5.

### Sex-specific DNA methylation alterations following pre-implantation alcohol exposure

We next sought to discern whether early pre-implantation alcohol exposure could have a sex-specific impact on later forebrain DNA methylation patterns. By analyzing male and female samples separately (min. 3 samples/condition/sex, ≥10x sequencing depth), we identified 83 424 and 126 857 unique sequenced tiles in male and female forebrains, respectively (**Fig. 4A**). Although there were lesser regions analyzed in males, they showed a larger number of DMRs following pre-implantation alcohol exposure (DMRs: males n=2 097, females n=1 273) (**Fig. S6**). In male EtOH-exposed forebrains, we found 2097 DMRs of which 1936 (92%) showed decreased methylation. Comparably, female EtOH-exposed forebrains revealed 1273 DMRs of which 1066 (84%) presented decreased methylation (**Fig. S6A-D**). We also observed contrasts between DMR number, as well as associated biological processes among sexes (**Fig. S6E**). However, we do not exclude that these results could be related to only a partial overlap in sequenced regions (n=46 475) between male and female forebrain samples (**Fig. 4A**).

**Figure 4.**
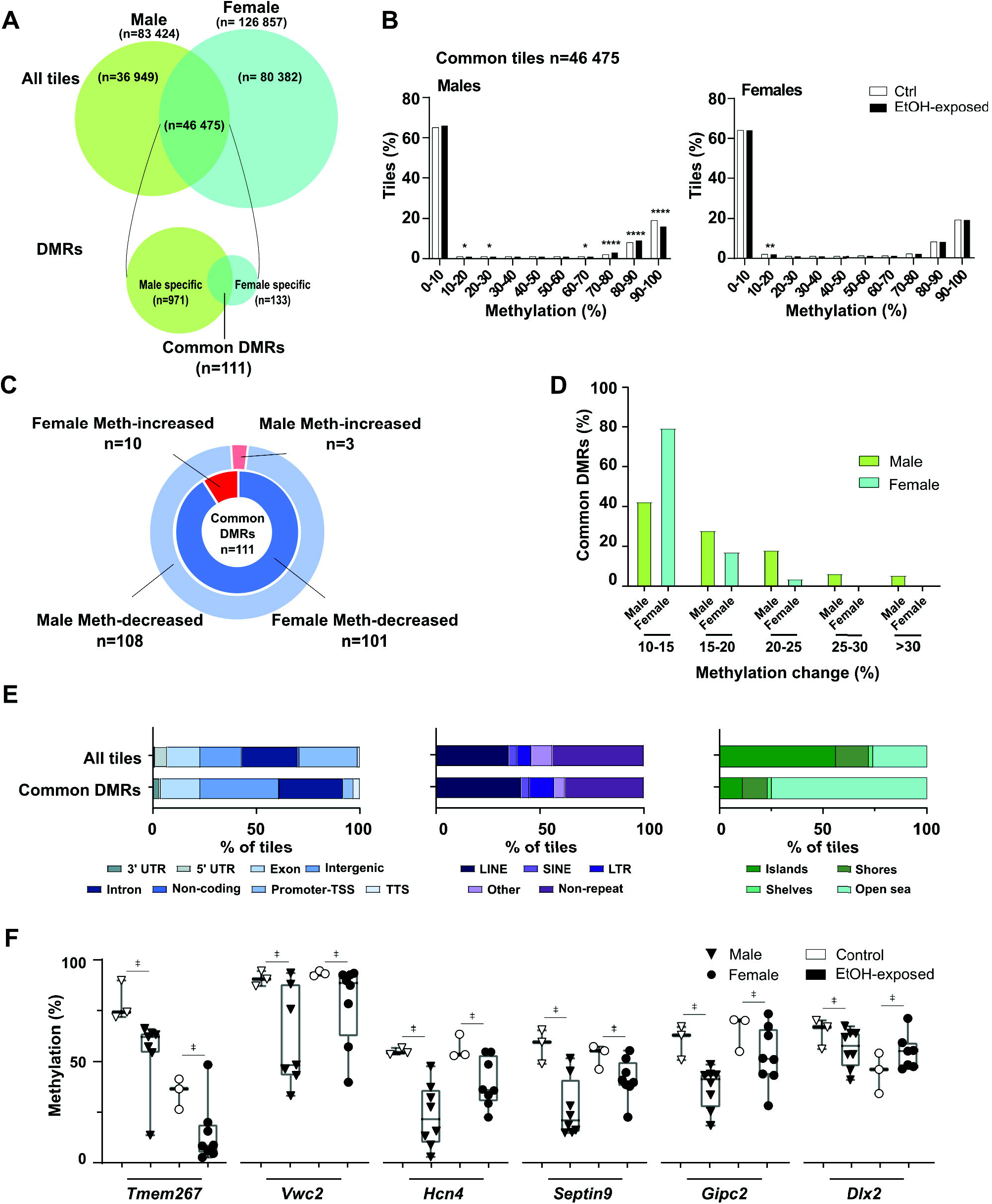
Sex-related changes in embryonic forebrain DNA methylation mediated by early pre-implantation ethanol exposure. **A)** Schematic design of sex-related genome-wide CpG methylation analysis in male (Ctrl n=3, EtOH n=8) and female (Ctrl n=3, EtOH n=8) E10.5 forebrain samples. Identification of all-tiles associated with either male samples (n=83 424), female samples (n=126 857), male-female common samples (n=46 475), as well as male-specific (n=971), female-specific (n=133) and common (n=144) DMRs (see methods section for details). **B)** Distribution of genomic tiles (100bp) (common tiles; n=46 475) across ranges (10%) of CpG methylation in male and female control and EtOH-exposed samples. ****p<0.0001, **p<0.01, *p<0.05; z-test proportion test. **C)** Distribution of common DMRs (n=111) with increased or decreased CpG methylation in male (outer circle) and female (inner circle) samples. **D)** Proportion of common DMRs associated with the changes of CpG methylation levels between control and EtOH-exposed in male and female forebrains. **E)** Percentage of tiles associated with various genomic features: genomic annotation (left), repeat elements (middle) and CpG-rich context (right) in common all-tiles (n=46 475) and common DMRs (n=111). **F)** Examples of CpG methylation levels of individual samples of common DMRs in male and female samples. □ represents significant differences in CpG methylation levels of DMRs (e.g., ±>10% methylation difference, q<0.01) between control and EtOH-exposed embryos (see methods section for details).

To circumvent this issue, we focused on the 46 475 regions with sufficient sequencing coverage in both sexes encompassing 373 530 CpGs (**Fig. 4A**). We did not observe any differences in global DNA methylation levels between sexes in either controls or ethanol-exposed forebrains (males: 29.4% vs 28.7%, females: 29.4% vs 29.2%; not shown). However, when these common tiles (n= 46 475) were distributed according to their DNA methylation levels in control and ethanol-exposed forebrains, we observed a greater shift in tile distribution in male EtOH-exposed samples (**Fig. 4B**), suggesting a greater effect on males. Accordingly, among the 46 475 male-female common tiles, we identified 971 male-specific DMRs, 133 female-specific DMRs, and 111 DMRs that were present in both sexes (**Fig 4A**). Out of these 111 common DMRs, 100 had similar alteration profiles (i.e., n=99 DMRs with decreased methylation, n=1 DMR with increased methylation) in male and female EtOH-exposed forebrains (**Fig. 4C**). For the other DMRs (n=11), the alcohol exposure caused conflicting alteration profiles between sexes (e.g., decreased-methylation in males vs increased-methylation in females). For the most part, the common DMRs displayed low levels of methylation changes (10-15% range) for both sexes (42% for males, 79% for females), with methylation changes greater than 25% only present in male EtOH-exposed samples (**Fig. 4D**). When compared to the 46 475 common tiles analyzed, the 111 common DMRs were more enriched in intergenic regions and intron categories, as well as for LINE elements, whereas they were mostly depleted of CpG rich sequences (i.e. CpG islands). In addition, common DMRs in male and female EtOH-exposed forebrains showed various levels of DNA methylation for an assortment of genes, including *Tmem267* (putative oncogene), *Vwc2* (neural development and function), *Hcn4* (cardiac function), *Septin9* (cytoskeletal formation), *Gipc2* (gastrointestinal processes) and *Dlx2* (forebrain and craniofacial development) (**Fig. 4F**, **Fig. S7**).

When we turned our attention to male-specific (n=971) and female-specific (n=133) DMRs within the 46 475 common tiles, we observed that pre-implantation alcohol exposure had a more profound impact on male forebrains in terms of DMR number and level of methylation changes (**Figs. 4A**, **5A**). Although a portion of observed DMRs showed increased methylation (males 3%; n=31, females 23%; n=30), the majority of sex-specific DMRs were associated with a partial loss of DNA methylation (males n=940, females n=103) in all chromosomes. The only chromosomes that showed an enrichment (p<0.0001) in DMRs were female X-chromosomes (n=48/133). Most of these sex-specific DMRs showed low levels of alterations, in the 10-15% range (males n=627 DMRs; 65%, females n=112 DMRs; 84%), with males having a larger proportion of DMRs with >15% methylation changes (**Fig. 5B**). When we highlighted genomic features associated to these sex-specific DMRs, we observed more female-specific DMRs in promoter regions (17% vs 4%) and CpG rich regions (33% vs 8%) when compared to male-specific DMRs (**Fig. 5C**). When we focused on the distribution of CpG sites in a sequence context (i.e., CpG islands, shores, shelves), we noticed a significant loss of global methylation in male-specific DMRs for male EtOH-exposed forebrains, and similarly for female-specific in female EtOH-exposed forebrains (**Fig. 5D**). For female-specific DMRs, we noticed that DNA methylation levels in control forebrains are all higher in females compared to males. These higher methylation levels are mainly associated with the presence of methylation marks associated to the X-inactivation process in females (female-specific DMRs on X-chromosome: CpG islands 35/44; shores 9/36, shelves 0/6. **Fig. 5F** bottom for examples). The sex-specific DMRs resulting from pre-implantation alcohol exposure were related to divergent biological processes between males (e.g., muscle differentiation, cell projection organization) and females (e.g., receptor protein tyrosine kinase pathway, epithelial cell migration) (**Figs. 5E, 5F**).

**Figure 5.**
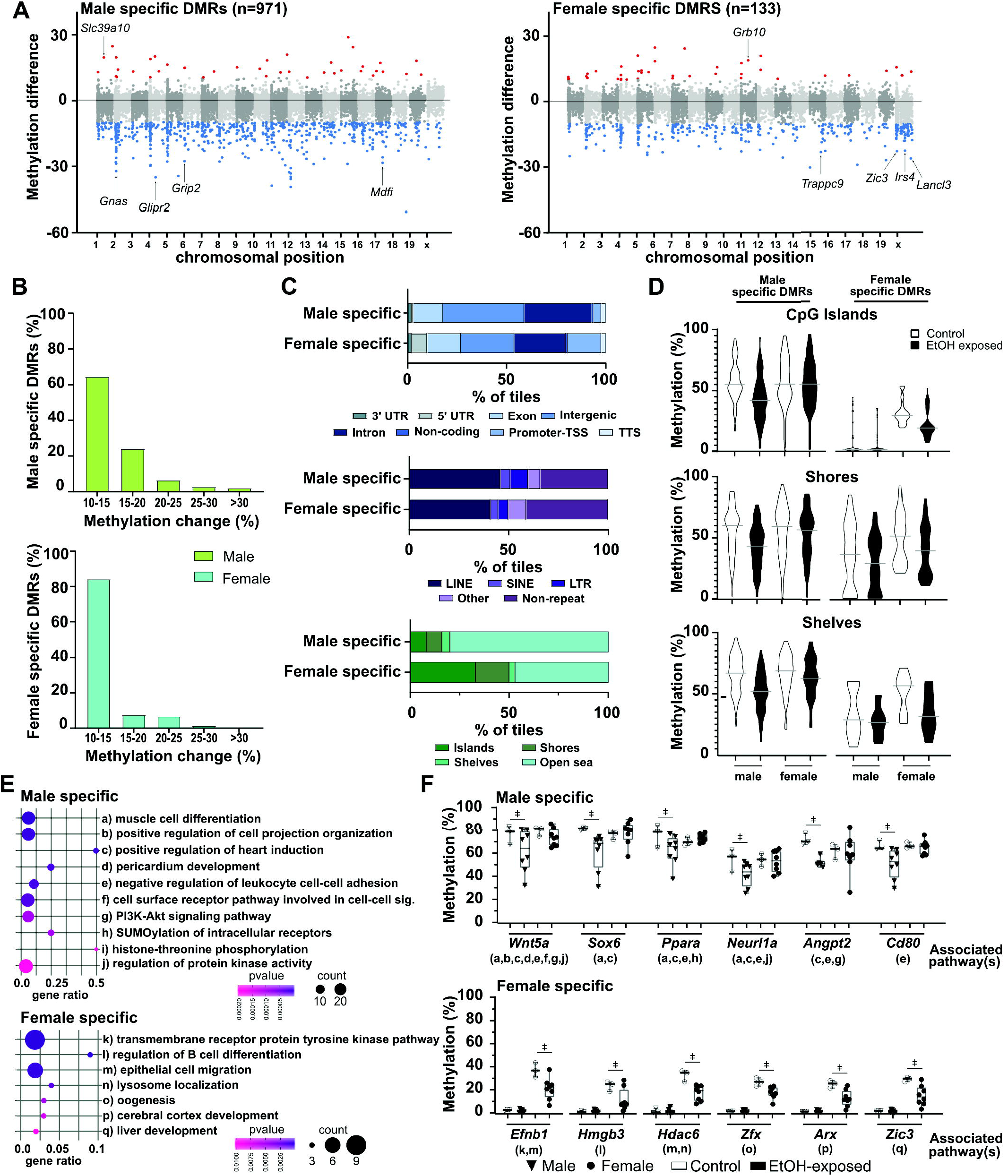
Early pre-implantation ethanol exposure induces sex-specific DNA methylation alterations in developing embryonic forebrain. Identification of male and female sex-specific DMRs in ethanol-exposed E10.5 forebrains is described in Fig.4A. **A)** Manhattan plot showing differences of CpG methylation by chromosomal position for male-specific (n=971; left graph) and female-specific (n=133; right graph) DMRs. Red dots represent DMRs with increased-methylation in EtOH-exposed forebrains (n=31; right graph and n=30; left graph); blue dots represent DMRs with decreased methylation in EtOH-exposed forebrains (n=940; right graph and n=103; left graph); grey dots represent tiles with less than 10% methylation differences between EtOH-exposed and control forebrains (n=45 504; right graph and n=46 342; left graph). **B)** Proportion of sex-specific DMRs (upper graph; male n=971, lower graph; female n=133) associated with changes in CpG methylation levels between control and EtOH-exposed E10.5 forebrains. **C)** Percentage of sex-specific DMR-associated tiles (male n=971, female n=133) across various genomic features: genomic annotation (top), repeat elements (middle) and CpG-rich context (bottom). **D)** Percentage of CpG methylation levels of male- and female-specific DMRs based on the distribution of CpG sites in CpG islands, CpG shores and CpG shelves. **E)** Functional enrichment analysis showing top enriched pathways for male-(top 10 pathways, n=508 unique gene DMRs) and female-specific DMRs (n=87 unique gene DMRs), based on Metascape analysis for pathways and p-value. The size of the dot represents the number of DMR-associated genes in pathways, and gene ratio represents the number of DMR-associated genes with regards to the number of genes in a pathway. **F)** Examples of CpG methylation levels of individual samples for sex-specific DMR-associated genes related to the top enriched pathways in E). Letters under gene name relate to the pathways in E). □ represents significant differences in CpG methylation levels of DMRs (e.g., ±>10% methylation difference, q<0.01) between control and EtOH-exposed embryos (see methods section for details).

We then evaluated whether early embryonic alcohol exposure led to expression errors in the forebrains of E10.5 embryos by focussing on a group of genes (with or without DMRs) implicated in the regulatory network coordinating the timing of GABAergic interneuron migration and forebrain formation. At the core of this network is the *Dlx* family of homeodomain transcription factors (50, 51). We show that *Dlx2* (DMR in gene body; males and females) and *Dlx1* (no DMRs) have small but significant changes in gene expression (*Dlx2*: males and females; *Dlx1*: females) (**Fig. S8A**), whereas for *Dlx5* and *Dlx6* (no DMRs), the expression remained unchanged (**Fig. S8B**). Upstream key regulator of MGE (medial ganglionic eminence)-derived GABAergic interneurons (52–54), *Nkx2.1* (no DMRs) did not show expression alterations (**Fig. S8B**), however downstream transcription factors such as *Sox6* (DMR in gene body; males), and *Arx* (direct target of *Dlx2* (55, 56)), DMR in promoter; females) showed small but significant gene expression alteration (*Sox6*; males and females, *Arx*; males) (**Fig. S8A**).

Together, these results indicate that alcohol exposure in pre-implantation embryos in conjunction with epigenetic reprogramming leads to variable levels of sex-specific alterations, with male embryonic forebrains being more prone to DNA methylation alterations. This suggests that pre-implantation male embryos are more susceptible to the initial adverse exposure rendering them less efficient at re-establishing proper DNA methylation patterns during the *de novo* methylation wave, or that female embryonic cells are better at rectifying dysregulated DNA methylation patterns during development.

### Pre-implantation alcohol exposure leads to partial loss of imprinted DNA methylation patterns

To further determine if a binge alcohol exposure episode on early embryos undergoing the epigenetic reprogramming wave is more adverse on the DNA methylation patterns of male or female embryos, we directed our attention to ICRs of imprinted genes, which are well known for their key roles in brain development and growth. We and others have shown that allele-specific methylation maintenance is required on ICRs during the reprogramming wave, as partial or complete loss of ICR profiles is permanent. Since our alcohol exposure specifically targets E2.5 embryos (8-cell stage), dysregulation in ICRs methylation maintenance would still be detectable in E10.5 forebrains. From the 46 475 commonly sequenced tiles, 28 were located within the ICRs of 9 imprinted genes. Our non-allele specific differential methylation analysis revealed that out of these 28 regions, 24 were differentially methylated in male EtOH-exposed forebrains. These 24 DMRs were associated to imprinted genes *H13*, *Nnat*, *Gnas*, *Kcnq1*, *Plagl1*, *Zrsr1*, *Peg13* and *Igf2r* (**Fig. 6A**). In females, only 2 of those 28 tiles showed altered levels in female EtOH-exposed forebrains, which were associated with *Gnas* and *Grb10*. We expanded our search for dysregulated ICR DNA methylation patterns within uniquely sequenced regions in male (n=36 949 tiles) and female (n=80 382 tiles) forebrains (**Fig. 4A**) to retrieve additional ICR-associated tiles. Again, in EtOH-exposed male forebrains, the majority of ICRs (10/13 tiles) showed altered DNA methylation patterns (**Fig. 6B**), which were associated with *Gnas*, *Snrpn*, *Peg3*, *Plagl1*, *Zrsr1* and *Impact.* Conversely, in EtOH-exposed female forebrains, we now observed a large portion of ICRs (19/36 tiles) with dysregulated DNA methylation levels associated to various imprinted genes (i.e., *Gnas*, *Peg10*, *Inpp5f*, *Kcnq1*, *Snrpn*, *Grb10*, *Zrsr1*, *Peg13*, *Slc38a4*, *Igf2r* and *Impact*). When we plotted individual sample DNA methylation values for these ICR-associated DMRs, we again observed a high degree of heterogeneity in EtOH-forebrains with some samples showing altered DNA methylation levels (e.g., partial loss, complete loss), and others revealing normal control methylation values (**Fig. 6A-C**).

**Figure 6.**
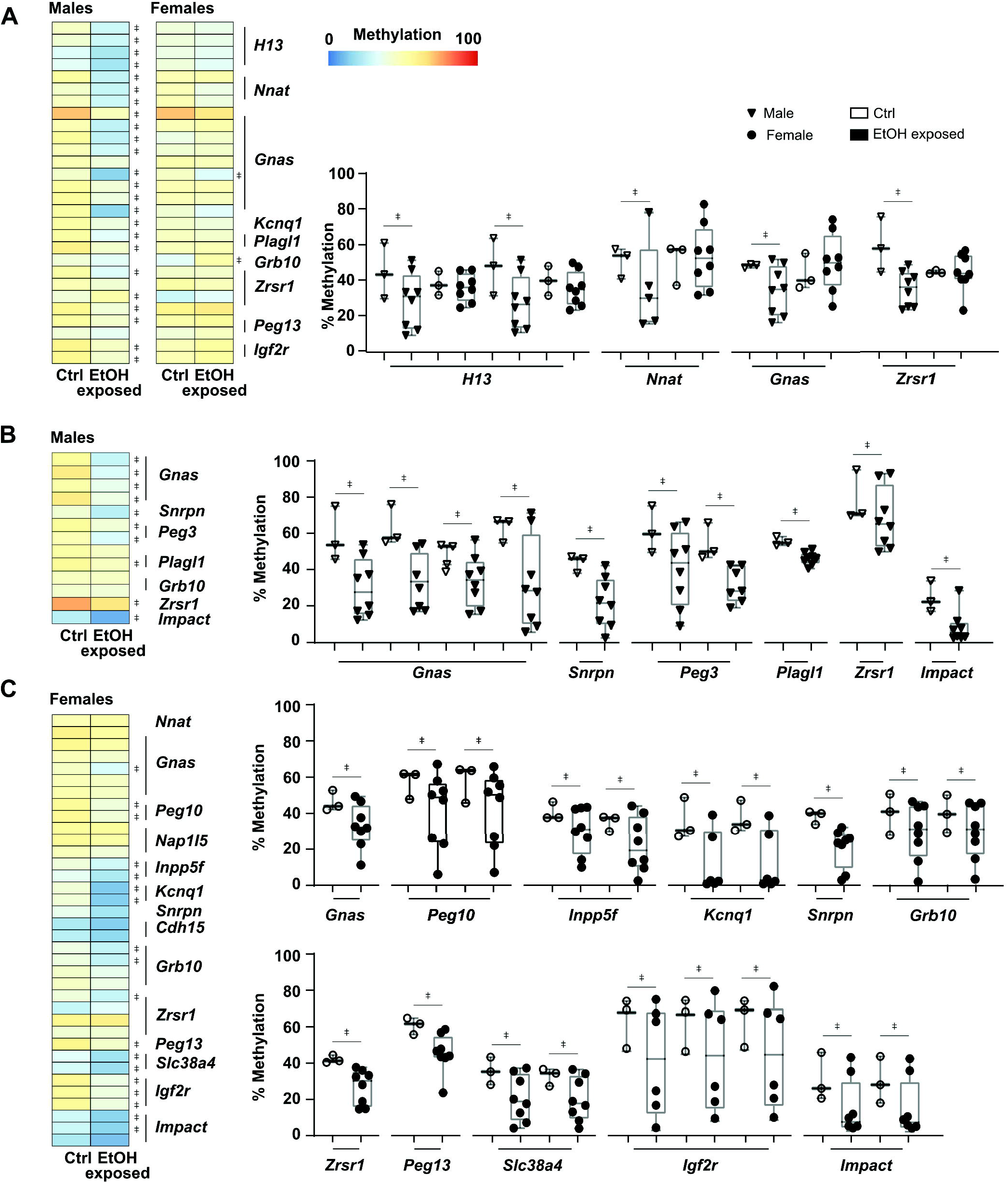
Early pre-implantation ethanol exposure leads to partial loss of DNA methylation maintenance across imprinting control regions. **A-C)** Heatmaps representing CpG methylation levels for control versus EtOH-exposed samples in regions (100bp tiles) located within defined imprinting control regions. Examples of CpG methylation levels of individual samples in imprinting control regions associated tiles are shown. □ represents significant differences in CpG methylation levels of DMRs (e.g., ±>10% methylation difference, q<0.01) between control and EtOH-exposed embryos (see methods section for details). **A)** Imprinting control regions (n=28 tiles) analyzed (sufficient sequencing coverage) in both male and female samples (control vs EtOH-exposed). **B)** Imprinting control regions (n=13 tiles) analyzed only in males (lack of proper sequencing coverage in females). **C)** Imprinting control regions (n=36 tiles) analyzed only in females (lack of proper sequencing coverage in males).

This partial loss of DNA methylation signatures within ICRs of male and female E10.5 EtOH-exposed forebrains was associated with small but significant deviation in expression level for a number of imprinted genes (e.g., *Gnas*, *Plagl1*, *Peg13*) (**Fig. S9A**). Whereas for others, although we observed similar ICR-associated DMRs, we did not detect any alteration in imprinted gene expression (e.g., *Impact*, *Peg10*, *Grb10*) for either male or female E10.5 EtOH-forebrains (**Fig. S9B**).

Both male and female EtOH-exposed forebrains showed variable levels of DNA methylation alterations in imprinted gene ICRs, which suggests that a binge-like alcohol exposure during pre-implantation interferes with the DNA methylation maintenance machinery during the epigenetic reprogramming wave. Finally, these results also suggest that male pre-implantation embryos are more susceptible to the initial adverse of alcohol exposure.

## DISCUSSION

In this study, we showed that a binge alcohol exposure episode on early-stage embryos (8-cell; E2.5) leads to a surge in morphological brain defects and delayed development during fetal life, that are reminiscent of clinical features associated to FASD. As seen in children exposed to alcohol prenatally, a portion of ethanol-exposed embryos presented a spectrum of alcohol-induced macroscopic defects while the majority showed no noticeable dysmorphic features and not alterations. However, forebrain tissues from ethanol-exposed embryos with no visible macroscopic abnormalities, developmental delays, alteration in cell proliferative response or cell death still presented lasting genome-wide DNA methylation alterations in genes associated to various biological pathways, including neural/brain development, and tissue and embryonic morphogenesis. These ethanol-exposed embryos also showed partial loss of imprinted DNA methylation patterns for various imprinted genes critical for fetal growth, development, and brain function. Moreover, we observed alcohol-induced sex-specific errors in DNA methylation patterns with male-embryos showing increased vulnerability.

### Modeling early pre-implantation alcohol exposure

One of the challenges when modeling FASD is unscrambling direct and indirect outcomes associated with the amount, pattern (continuous *vs.* binge drinking), and developmental timing of alcohol exposure (18, 19, 45, 57–63). By targeting pre-implantation embryo, we observed that a binge-like alcohol exposure on a single embryonic cell type (8-cell stage blastomeres) leads to increased rate of macroscopic defects (e.g., brain anomalies, growth restriction, heart defects) across litters, with no impact on litter size or on sex-specific phenotypic representation during fetal life. These findings corroborate with pioneer work that reported abnormal fetal development without significant reduction in litter size following exposure during pre-implantation (64, 65). However, the acute dosage regimen paradigms used in those studies resulted in a much higher rate (67% to 100%) of embryonic abnormalities and severe growth retardation, as well as fetal death (41, 66). Nevertheless, such studies along with others (67) confirm that even prior to direct maternal-fetal interface exchanges via the placenta, alcohol can reach the developing pre-implantation embryos through the female reproductive track. *In vitro* studies support that pre-implantation embryos are sensitive and negatively affected by alcohol exposure (68). Although less investigated and understood than maternal exposure, studies suggest that alterations (e.g., epigenetic errors) initiated on the fathers’ sperm are passed-on during fertilization to pre-implantation embryos, influence development beyond implantation, and lead to abnormal offspring development (e.g., fetal growth restriction, birth defects, placental defects) (58, 69–72).

It remains to be defined whether some of the milder abnormalities or delays that we observed at mid-gestation would become resolved or accentuated by birth, and whether embryos that presented no visible abnormalities or developmental delays but had DNA methylation alterations would show cognitive dysfunctions as observed in other FASD-models and children with FASD. In parallel, outlining if the dysregulation in DNA methylation profiles is associated with abnormal migration and organization of specific brain cell subtypes during development would further extend our understanding about the effect of early pre-implantation alcohol exposure on the neurobiological phenotype in offspring.

### Pre-implantation alcohol exposure leads to partial loss of imprinted DNA methylation

Our genome-wide, high-resolution analysis of ethanol-induced DNA methylation alterations highlighted partial loss of DNA methylation maintenance within various 100bp tiles located across imprinting control regions (e.g., *Gnas*, *Zrsr1, Impact*) in E10.5 male and female forebrains. Then again, methylation alterations were not necessarily widespread across entire imprinting control regions, or in all individual ethanol-exposed embryos, suggesting only a slight reduction in DNMT1/DNMT1o maintenance activity. Concordantly, some imprinted genes showed alterations in their expression profiles (e.g., *Gnas*, *Plagl1*, *Peg13*) in both male and female EtOH-exposed forebrains, whereases others (e.g., *Impact*, *Peg10*, *Grb10*) were not affected. We know that temporary lack of DNA methylation maintenance by DNMT1o in 8-cell embryos (E2.5) leads to delays in development and to a wide range of lethal anatomical abnormalities that are associated with epigenetically mosaic embryos that failed to properly maintain normal imprinted methylation patterns at mid-gestation (73–75). In zebrafish, alcohol exposure during the two first days of embryo development led to reduced levels of *Dnmt1* expression (76), which could ultimately lead to a temporary reduction in methylation maintenance activity. Studies aiming at defining how pre-implantation embryos respond to environmental stimuli (e.g., assisted reproductive technologies, toxicants, ethanol) have broadly explored the impact on imprinted genes, but have mainly relied on evaluating DNA methylation levels of imprinting control regions using targeted approaches (i.e., specific genomic loci) (41, 77, 78). Although informative, conclusions are often based on profiling DNA methylation levels of a limited number of CpG sites associated to a handful of genes. For instance, modeling binge alcohol exposure during two consecutive days (E1.5, E2.5) led to decreased fetal and placental weight by E10.5, but only to a partial loss of DNA methylation in the *H19* —a negative regulator of growth and proliferation— imprinting control region (only 17 CpG analysed) in the placenta. Although only a portion of one ICR was evaluated, these results insinuated that embryonic imprinted methylation was not affected by such early pre-implantation alcohol exposure.

Since imprinted genes are well recognized in regulating essential neurodevelopmental processes, including neural differentiation, migration and cell survival (79), we can presume that the forebrain of ethanol-exposed embryos with severe macroscopic defects and developmental delays would have broader loss of imprinted DNA methylation patterns compared to those with no visible abnormalities. However, the fact that morphologically normal forebrain tissues derived from ethanol-exposed pre-implantation embryos reveal widespread low-level DNA methylation alterations with increased inter-individual variability in imprinted control regions supports that failure in maintaining accurate methylation imprints could contribute to the invisible nature of FASD.

### Pre-implantation alcohol exposure initiates sex-specific DNA methylation programming errors

We demonstrated that exposing 8-cell embryos to alcohol is detrimental for future forebrain autosomal and X-chromosome DNA methylation patterns. Furthermore, such an early embryonic exposure leads to sex-specific DNA methylation alterations with male embryos being more susceptible to alterations. Early embryonic developmental stages are marked by a series of molecular events that are crucial for the proper establishment of the developmental program, as well as the *de novo* genome-wide DNA methylation signatures that are conserved throughout development (29, 80). A considerable amount of evidence shows that interfering with this process *in vivo* or *in vitro* leads to sex-specific long-term effects in the offspring (36, 72, 81–87). Although the mechanism remains unclear, we know that molecular events in early-stage embryos are marked with sex-specific discrepancies. For instance, mammalian pre-implantation embryos display differential expression of sex chromosome and autosomal transcripts, leading to extensive transcriptional sexual dimorphism (88, 89). In mouse, sex-biased gene expression (n=69; mainly X-linked) is detected as early as the 8-cell stage with substantial variation between individual embryos, and with female embryos expressing higher transcript levels. Then, just prior to the *de novo* re-methylation wave, the sex chromosomes seem to further drive this sexual dimorphism in transcriptional regulation for hundreds of autosomal genes, with *Dnmt3a* and *Dnmt3b* showing higher expression in male blastocyst (90). This is consistent with sex-specific acquisition of DNA methylation reported in bovine blastocyst, with males showing increased levels (89, 91). Thus, the alcohol-induced sex-specific DNA methylation alterations resulting from an early embryonic alcohol exposure model are mostly initiated at the 8-cell stage because of the male-female differences in transcriptional regulation. The DNA methylation alterations would then trigger a series of events that would negatively impact the re-establishment of genome-wide *de novo* DNA methylation profiles occurring between E3.5 and E6.5 in a sex-specific manner, with males being more affected. Once established, these abnormal *de novo* DNA methylation profiles would be maintained or would initiate further DNA methylation alterations during subsequent developmental stages. Since the sex-specific alterations observed in the forebrains are linked to divergent biological pathways, this could lead to sex-specific neurocognitive impairments in offspring. Although such observations have been made in children with FASD, with males appearing to be more vulnerable to the irreversible effects of fetal alcohol exposure (e.g., cranial and facial malformations, learning disabilities, social and memory disabilities, and altered brain structure and function) (reviewed in (92)), further studies are needed to explore the long-term degree and impact of the sexual dimorphism in this pre-implantation alcohol exposure paradigm.

### Pre-implantation alcohol exposure and impact on neurodevelopment

One common phenotype observed in human FASD and animal models of prenatal alcohol exposure is the interference of alcohol on the development of the central nervous system (structural or functional abnormalities). We observed DNA methylation alterations in promoters and bodies of genes implicated in brain and nervous system function and development (e.g., *Dlx2, Epha7*, *Foxa1, Nkx6, Vwc2*, *Sox6*). For example, *Dlx2* (Distal-Less Homeobox 2), is part of a transcription factor family (*Dlx1/2*; *Dlx3/4*; *Dlx5/6*) that is critical molecular determinants for forebrain and craniofacial development, as well as for coordinating the timing of GABA(gamma-aminobutyric acid)ergic interneuron migration and process formation (93–96). *Dlx2* is required to promote the expression of several downstream factors including other *Dlx* genes and *Arx* (directly activated by *Dlx2*) *(56)*, an X-linked gene that also controls cortical interneuron migration and differentiation (97), which incidentally showed altered DNA methylation in female ethanol-exposed forebrains. Ethanol-exposed forebrains showed reduced methylation in *Dlx2* (gene body) and *Arx* (promoter), which correlated with reduced *Dlx2* and *Arx* expression, whereas the average expression of *Dlx5* and *Dlx6* (no DMRs) remained unchanged. Mice lacking *Dlx1/2* have profound deficits in tangential migration of GABAergic cortical interneurons and neurite growth. Similarly, prenatal stress-induced anxiety in mouse leads to GABAergic interneuron deficiency-associated dysregulation of promoter DNA methylation levels of *Gad67* (glutamic acid decarboxylase 67), an enzyme critical for GABA synthesis (the principal inhibitory neurotransmitter). In prenatal alcohol exposure models targeting different brain developmental time points, the migration and positioning of GABAergic cortical interneurons are profoundly impaired, leading to subsequent cortical dysfunction (98–102). We know that cortical interneuron dysfunction, associated with impaired development, migration, or function of interneurons, results in interneuronopathies that contribute to multiple neurodevelopmental disorders including autism, epilepsy, schizophrenia, and FASD (103–105). We also know that GABAergic interneurons are particularly responsive to adverse maternal exposures during *in utero* periods of developmental plasticity; from the moment these cells arise, to the shaping of cortical circuits (101, 105). Our data suggest that early embryonic alcohol exposure triggers alterations in the epigenome that leave lasting signals that could lead to pathological plasticity in the developing brain. We need to further define whether GABAergic interneurons, or other cortical neurons, are particularly vulnerable to early embryonic alcohol-induced epigenetic programming errors, and whether this could drive interneuronopathies or neurodevelopmental impairments associated with FASD.

### Study Limitation

One limitation to our study is the absence of information about the dose-dependency of alcohol exposure with regards to the phenotypes and molecular consequences observed. Since higher peak blood alcohol concentrations have been shown to play a critical role in the extent of prenatal alcohol exposure-related damages (17, 18, 45, 48, 66, 86, 106), we can presume that similar observations would be observed using our paradigm. Nonetheless, the impact of low-dose and early prenatal alcohol exposure should not be overlooked, as they have been associated to increase the risk of mental illness regardless of a FAS or FASD diagnosis in human (review in (107)). To fully comprehend the wide range of detrimental consequences (visible and invisible) associated to dose-dependency of alcohol exposure during pre-implantation will require a thorough investigation across pre- and post-natal development.

## CONCLUSION

We showed that pre-implantation alcohol exposure is detrimental for normal development and leads to a broad spectrum of adverse outcomes that closely replicate clinical facets observed in children with FASD. Specifically, we demonstrated that a binge-like drinking episode while pre-implantation embryos are in the mist of their reprogramming wave leads to two main categories of lasting programming errors: the partial loss of DNA methylation at several imprinted control regions, and the abnormal re-establishment of *de novo* DNA methylation profiles in key biological pathways (e.g., neural/brain development, tissue and embryonic morphogenesis). Further studies in peri-implantation embryos, when DNA methylation is globally reacquired, and in specific cell subtypes during key neocortex developmental time points will provide a better understanding of the fundamental mechanisms leading to these DNA methylation programming errors and their implication in neurodevelopmental FASD-related deficits. Importantly, our data demonstrate that pre-implantation alcohol exposure does not lead to an “all-or-nothing” response, as morphologically normal embryos still presented conserved and sex-specific DNA methylation alterations in forebrain tissues, which could be indicative of the particular sexual dimorphism in cognitive dysfunctions associated with FASD. Thus, our study provides strong scientific evidence to refute the “all-or-nothing’’ principle and supports the potential contribution of early embryonic epigenetic alterations to the manifestation of neurodevelopmental phenotypes observed in a portion of children with FASD.

## METHODS

### Pre-implantation embryo binge-like ethanol exposure model

Animal work was approved by the Comité Institutionnel de Bonnes Pratiques Animales en Recherche (CIBPAR) of the CHU Ste-Justine Research Center under the guidance of the Canadian Council on Animal Care (CCAC). Female C57BL/6 mice (8-week-old) were mated with same age C57BL/6 males (Charles River laboratories). Females that showed copulatory plugs the next morning were considered pregnant with day 0.5 embryos (E0.5). They were separated from the males and housed together in a 12h light/dark cycle with unlimited access to food and water.

Using a recognized prenatal binge-like alcohol exposure paradigm (41, 45, 66, 106), pregnant females (E2.5) were injected with 2 subsequent doses of 2.5g/kg ethanol 50% (ethanol-exposed group) or an equivalent volume of saline (control group) at 2 hours intervals. Female with the same treatment were housed together and had negligible handling during the gestation.

### Blood alcohol concentration quantification

Blood alcohol concentration associated with our pre-implantation binge-like alcohol exposure paradigm was quantified over a 4h period. To avoid supplementary stress on experimental animals, a different subset of pregnant E2.5 females (n=12) was used for this experiment. Ethanol-exposed females (n=3) were euthanized at each time point (1, 2, 3 and 4h), blood was collected, samples were centrifuged to separate serum and plasma, and alcohol was quantified using the EnzyChrome Ethanol Assay kit (BioAssays Systems/Cedarlane) following manufacturer’s recommendations (1:5 plasma dilution).

### Morphological analysis

At E10.5, pregnant females were euthanized, and embryos were collected and dissected for morphological evaluation using Leica stereo microscope. Using LasX software, measurements of the crown-rump length (top of the head to the end of the tail), occipital to nose length (occipital part of the head to the nasal process), height of the head (top of the head to the beginning of the torso) and length of the midbrain (occipital part to the midbrain/forebrain limit) were done. Similarly, morphological defects (e.g., severe developmental delay or growth restriction, brain or head malformation, heart anomaly or any other unexpected feature) were evaluated (108–110). Three embryos (n=3) revealed more than 2 morphological defects and were placed in the category of their main defect (delayed n=2, brain malformation n=1). Embryos that hatched (Ctrl n= 23; EtOH n=21) from the yolk sac during dissection were excluded of both measurements and morphological analyses due to the possible deformation induced by the pressure on the embryo during the expulsion. The sex of each embryo was determined by qPCR using expression of *Ddx3*, on yolk sac DNA. Statistical analyses were done using GraphPad prism (version 8.4.3) for t-test with Welch correction and f-test for variance or R (version 3.5.0) for chi-square and proportional z-test.

### Histological analysis and immunostaining

Following collection, whole E10.5 embryos were fixed in 4% paraformaldehyde (PFA), post-fixed in EtOH 70% for 48h and paraffin embedded (111). Whole embryos were sectioned at 5μM and corresponding sections were stained with hematoxylin and eosin (112) to determine gross morphology, or for Cleaved Caspase-3 (Cell Signaling #9579S) or Ki67 (Abcam # AB15580) following manufacturer’s protocol and counterstained with hematoxylin. Imaging was done using Zeiss Zen Axioscan Slide Scanner system. Images processing and quantification were performed using ImageJ. Three similar sub regions of forebrain and midbrain (0.02mm^2^ each) were quantified across all samples. Statistical analyses were done using GraphPad prism (version 8.4.3) for t-test with Welch correction.

### DNA extraction and Reduced Representation Bisulfite Sequencing

Following morphological analyses, embryonic E10.5 forebrains were isolated, flash frozen and kept at −80°C. Genomic DNA was extracted from forebrains using the QIAamp DNA Micro kit (Qiagen #56304) following manufacturer’s recommendations. Extracted DNA was quantified using QuBit fluorimeter apparel with the Broad range DNA assay kit (ThermoFisher #Q32853). The sex of each DNA sample was again validated by *Ddx3* qPCR. DNA samples from forebrains with no apparent morphological defects were randomly selected, using 6 control embryos (3 males and 3 females, from 3 different litters) and 16 ethanol-exposed embryos (8 males and 8 females, from 6 different litters). EtOH-exposed and control embryo groups were similar in size and morphology (**Fig. S2**). Genomic DNA was used to produce rapid Reduced Representation Bisulfite Sequencing (rRRBS) libraries as previously described (27, 113–116). Briefly, 500ng of DNA was digested with *Msp1* restriction enzyme, adapters were attached to DNA fragments followed by sodium bisulfite conversion and amplification/indexation of libraries. Libraries were quantified using QuBit fluorimeter apparel with the High Sensitivity DNA assay kit (ThermoFisher #Q32854). Quality control was assessed using BioAnalyzer and paired-end sequencing was done on Illumina HiSeq 2500 at the Genome Québec core facility. We obtained between 19M and 41M raw reads for each sample (**Table S1**).

### Bioinformatics analysis

Data processing, alignment (mm10 genome) and methylation calls were performed using our established pipeline (27, 114–116) which includes tools such as Trim Galore (version 0.3.3) (117), BSMAP (version 2.90) (118) and R (version 3.5.0) (**Table S1**). Differentially methylated regions were obtained with MethylKit (version 1.8.1) (119) using the Benjamini-Hochberg false discovery rate (FDR) procedure. Fixed parameters were used, including 100bp stepwise tiling windows, a minimum of 2 CpGs per tile and a threshold of *q*<0.01. DNA methylation level is calculated as the average methylation of all CpGs within a tile for all the samples within a condition (**Fig. 3**: minimum 5 samples/condition, ≥10x sequencing depth; **Fig. 4–6** & **Fig. S6-S7**: minimum 3 samples/condition/sex, ≥10x sequencing depth). The number of CpGs per tile and bisulfite conversion rate (>97%) were obtained using a custom Perl script.

Annotation of the analysed tiles was done using Homer (version 4.10.1) with mm10 reference genome. Gene ontology enrichment analysis was performed with Metascape (120) online tool using only differentially methylated tiles located in genic regions. Repeats and CpG islands coordinates were obtained from UCSC table browser databases (mm10 genome). CpG context tracks (CpG shores and CpG shelves) were built by adding up respectively 0-2kb and 2-4kb from the CpG islands coordinates as previously describes (114, 121).

### RNA extraction and expression analysis by quantitative PCR (qPCR)

Quantitative gene expression analyses were performed as previously (122, 123). Briefly, embryonic E10.5 forebrains were isolated, flash frozen and kept at −80°C until RNA extraction. RNA was extracted using RNeasy Mini kit (Qiagen #74004) following manufacturer’s recommendations. Extracted RNA was quantified using QuBit fluorimeter apparel with the High Sensitivity RNA assay kit (ThermoFisher #Q32852). 600ng of RNA was used for cDNA conversion using SuperScript IV Reverse Transcriptase (ThermoFisher #18090010). For each gene (primer sequence **Table S2**), qPCR reactions were performed in triplicate on 5ng of cDNA using SensiFAST SYBR No-ROX (Bioline #BIO-98005) on a LightCycler 96 (Roche Life Science). Gene expression analysis and normalization was done using the 2^−ΔΔCt^ method using *Hprt1* and *Pgk1* as reference genes. Statistical analyses were done using GraphPad prism (version 8.4.3) for t-test with Welch correction.

## DATA AVAILABILITY AND MATERIALS

The data from this study have been submitted to the Gene Expression Omnibus (GSE162765). Reviewer link: https://www.ncbi.nlm.nih.gov/geo/query/acc.cgi?acc=GSE162765

## Token

exyxmucsxlilhwp

## ACKNOWLEDGMENTS

We thank the McGraw lab for critical comments and suggestions, as well as Elizabeth Maurice-Elder for editing.

## AUTHOR INFORMATION

### Affiliations

CHU Sainte-Justine Research Center, 3175 Chemin de la Côte-Sainte-Catherine, Montréal, QC H3T 1C5, Canada

Lisa-Marie Legault, Karine Doiron, Mélanie Breton Larrivée, Alexandra Langford-Avelar, Anthony Lemieux, Maxime Caron, Daniel Sinnett, Serge McGraw

Department of Biochemistry and Molecular Medicine, Université de Montréal, 2900 Boulevard Edouard-Montpetit, Montréal, QC H3T 1J4, Canada.

Lisa-Marie Legault, Mélanie Breton Larrivée, Alexandra Langford-Avelar, Anthony Lemieux, Serge McGraw

McGill University Health Centre Glen Site, 1001 Boulevard Décarie, Montréal, QC H4A 3J1, Canada.

Loydie Jerome-Majewska

Department of Pediatrics, McGill University, 1001 Boulevard Décarie, Montréal, QC H4A 3J1, Canada.

Loydie Jerome-Majewska

Department of Pediatrics, Université de Montréal, 2900 Boulevard Edouard-Montpetit, Montréal, QC H3T 1J4, Canada.

Daniel Sinnett

Department of Obstetrics and Gynecology, Université de Montréal, 2900 Boulevard Edouard-Montpetit, Montréal, QC H3T 1J4, Canada.

Serge McGraw

### Contributions

LML and SM conceptualized the study. LML and MBL contributed in data acquisition. LML, KD, ALA, AL, MC, LJM and DS participated in data analysis. LML, KD and SM wrote the manuscript. All authors read and approved the final manuscript.

### Corresponding author

Correspondence to Serge McGraw

## ETHICS DECLARATION

### Consent for publication

Not applicable

