## Supplemental Figures for "Pre-Implantation Alcohol Exposure Induces Lasting Sex-Specific DNA Methylation Programming Errors in the Developing Forebrain"

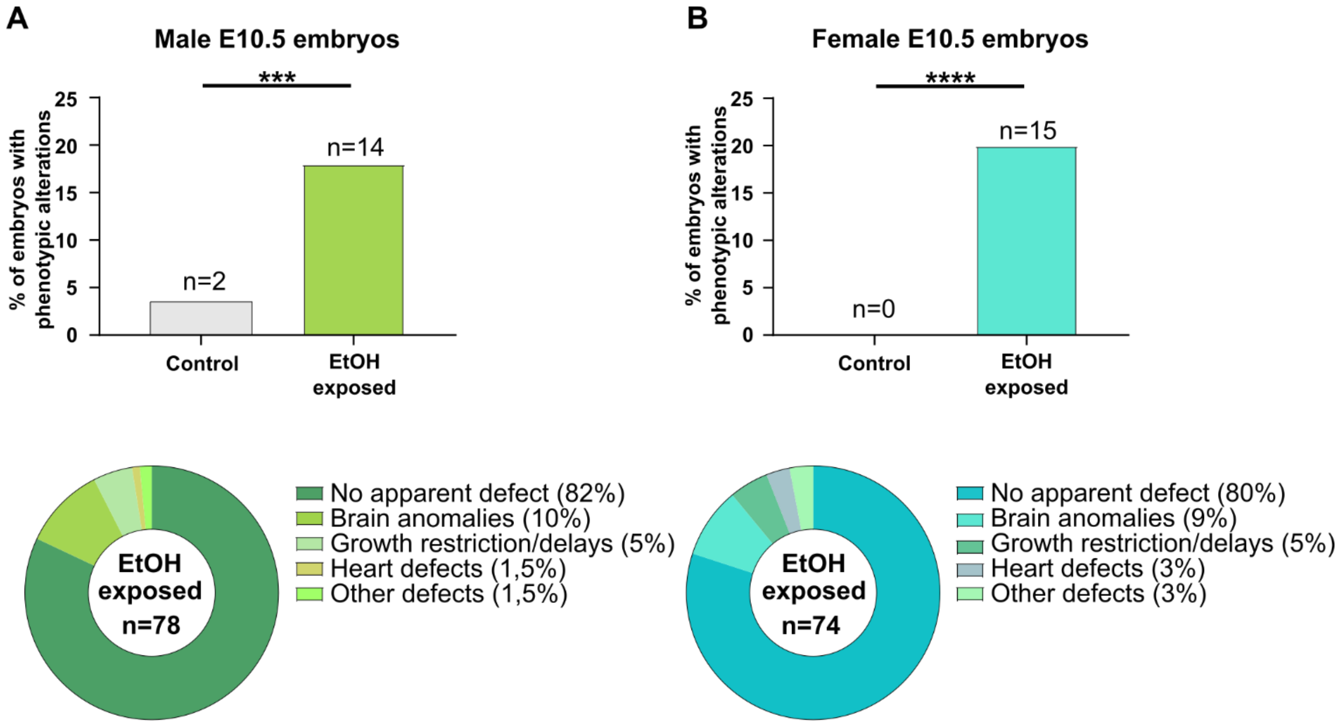


| | Ctrl | EtOH exposed | $x^2$ value | p-value |
|---|---|---|---|---|
| Embryos with phenotypic alterations | 2/55 | 14/78 | 13.73 | <0.001 |
| Mothers with at least one altered embryo | 2/16 | 10/22 | 4.66 | <0.01 |

| | Ctrl | EtOH exposed | $x^2$ value | p-value |
|---|---|---|---|---|
| Embryos with phenotypic alterations | 0/53 | 15/74 | 12.61 | <0.001 |
| Mothers with at least one altered embryo | 0/16 | 13/22 | 6.17 | <0.01 |

**Figure S1. No difference in phenotypic alterations between male and female developing embryos following early pre-implantation alcohol exposure**. **A)** Percentage of E10.5 male embryos with phenotypic alterations (Ctrl: 4%; n=2/55, EtOH-exposed: 18%; n=14/78, ***p<0.001; chi-square test) (upper graph). Classification and proportion of phenotypic alterations observed in EtOH-exposed male embryos (middle graph). Number of male embryos with alterations, and number of litters with at least one affected embryo. **p<0.01; chi-square test (lower table). **B**) Percentage of E10.5 female embryos with phenotypic alterations (Ctrl: 0%; n=0/53, EtOH-exposed: 20%; n=15/75, ***p<0.001; chi-square test) (upper graph). Classification and proportion of phenotypic alterations observed in EtOH-exposed female embryos (middle graph). Number of female embryos with alterations, and number of litters with at least one affected embryo. *p<0.05; chi-square test (lower table).

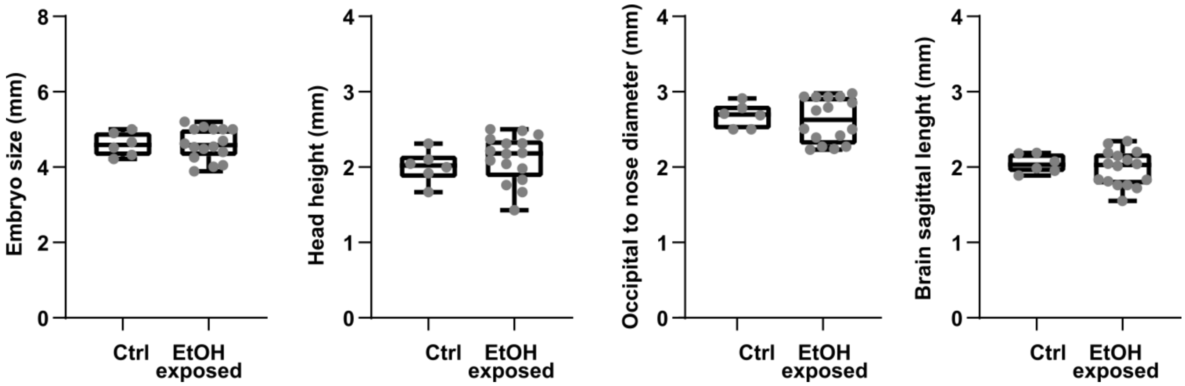


**Figure S2. Embryonic measurements of E10.5 embryos used to perform Rapid Reduced Representation Bisulfite Sequencing (rRRBS) on forebrains.** Crown-rump length, head height, occipital-nose length and brain sagittal length. Control embryos: (n=6, 3 litters), EtOH-exposed embryos (n=16, 6 litters). No significant difference; t-test with Welch's correction.

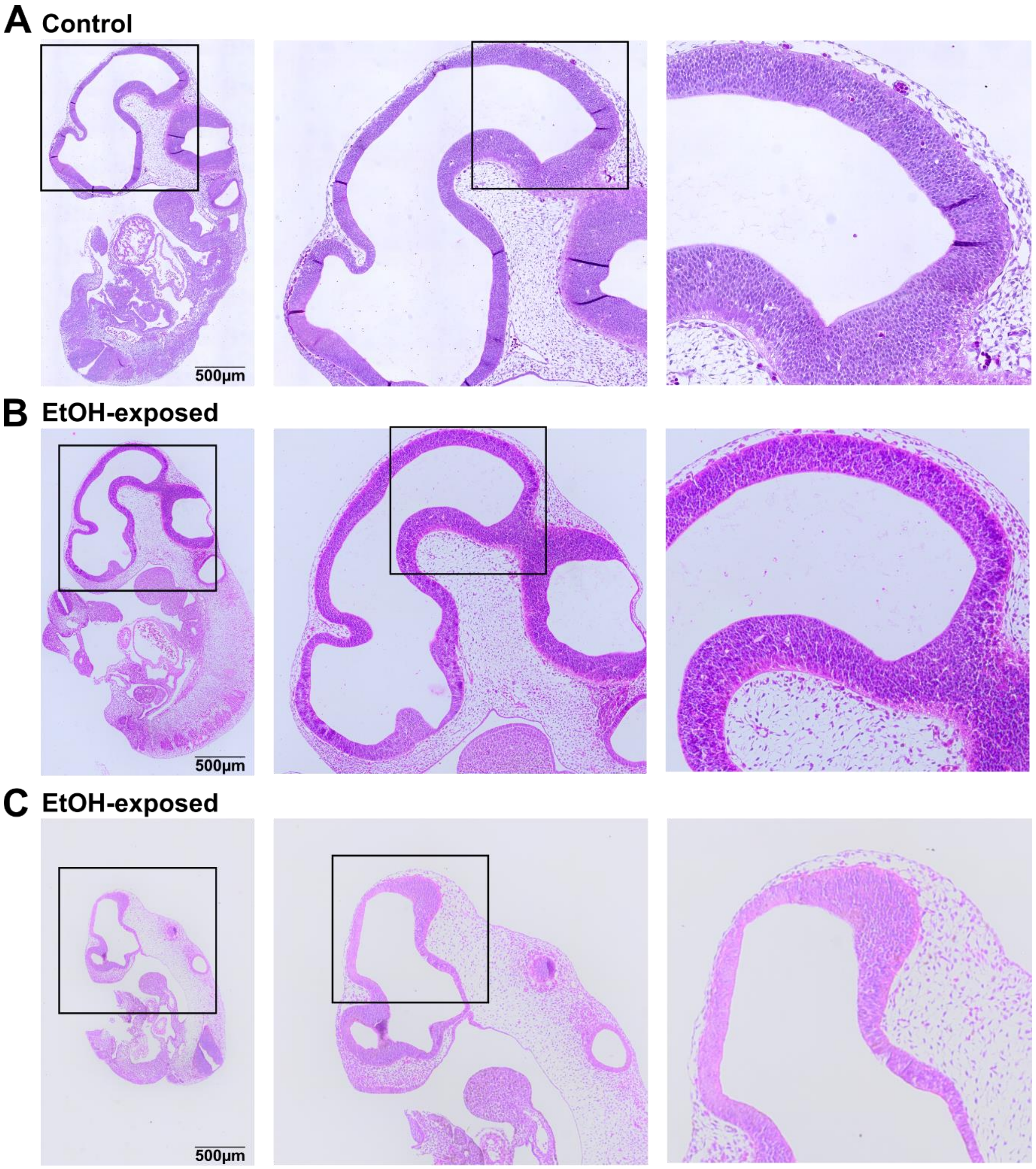


**Figure S3. Early pre-implantation alcohol exposure does not alter overall brain cell structural organization in mid-gestation embryos**. Hematoxylin and eosin (H&E) staining in control and ethanol-exposed E10.5 embryos (Male: ctrl n= 9, EtOH-exposed n=11; Female: ctrl n= 9, EtOH-exposed n=11). Representative images of H&E-stained sections of (A) control and (B, C) ethanol-exposed embryos, with focus on head area and sub-section of brain. A) Control male embryo. B) Male ethanol-exposed embryo without any morphological defect observed during dissection. C) Female ethanol-exposed embryo with a delayed development.

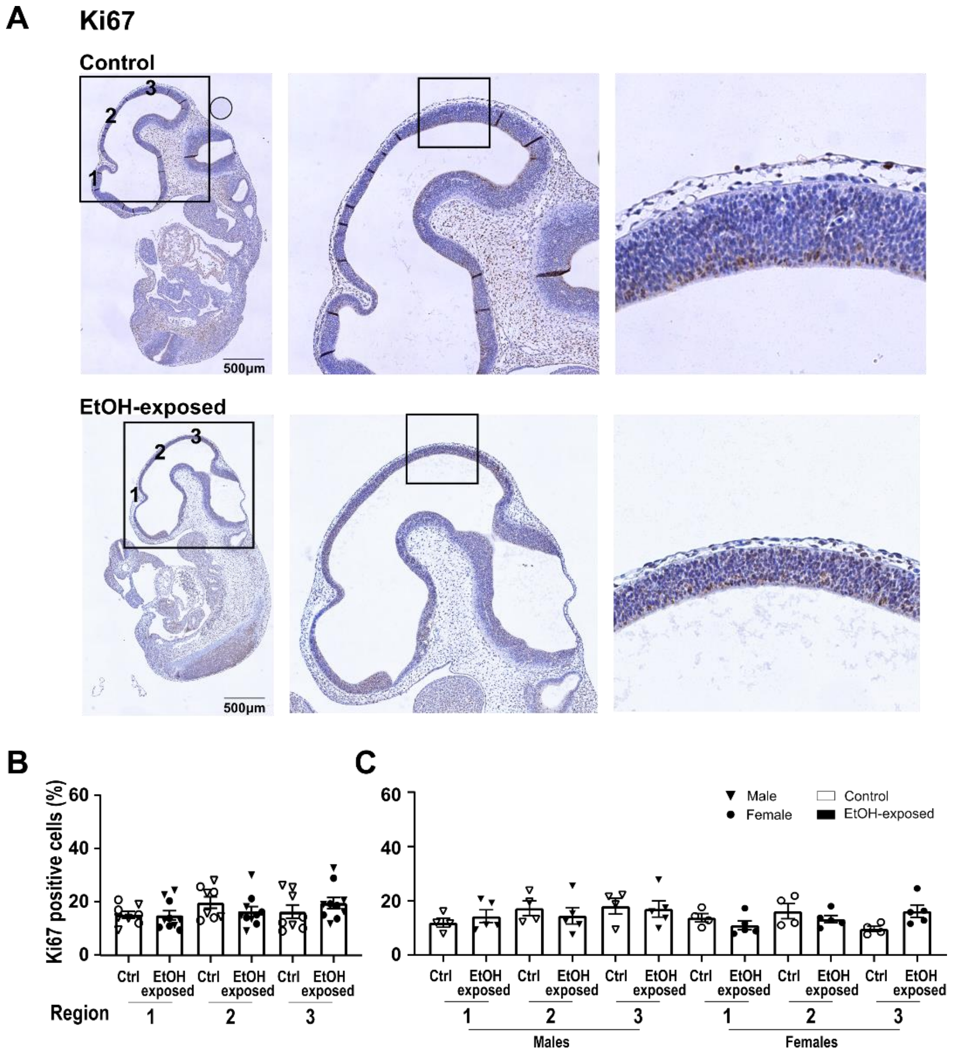


**Figure S4. Early pre-implantation alcohol exposure does not modify brain cell proliferative state in mid-gestation embryos**. Immunohistochemical staining for Ki67 (brown) and counterstaining with hematoxylin (blue) in control and ethanol-exposed embryos. A) Representative distribution of Ki67 positive cells in sagittal sections of E10.5 embryos. Left panel shows the three brain regions used to quantify Ki67 positive cells. Details of framed areas are shown in middle and right panels. B) Percentage (%) of Ki67 positive cells across brain regions in control and ethanol-exposed (no apparent morphological defects) embryos (males and females). Bars represent mean ± SEM. C) Sex-specific quantification of Ki67 positive cells. Data from B) separated in male and female samples; male (Ctrl n= 4, EtOH-exposed n=5), female (Ctrl n= 4, EtOH-exposed n=5). Bars represent mean ± SEM.

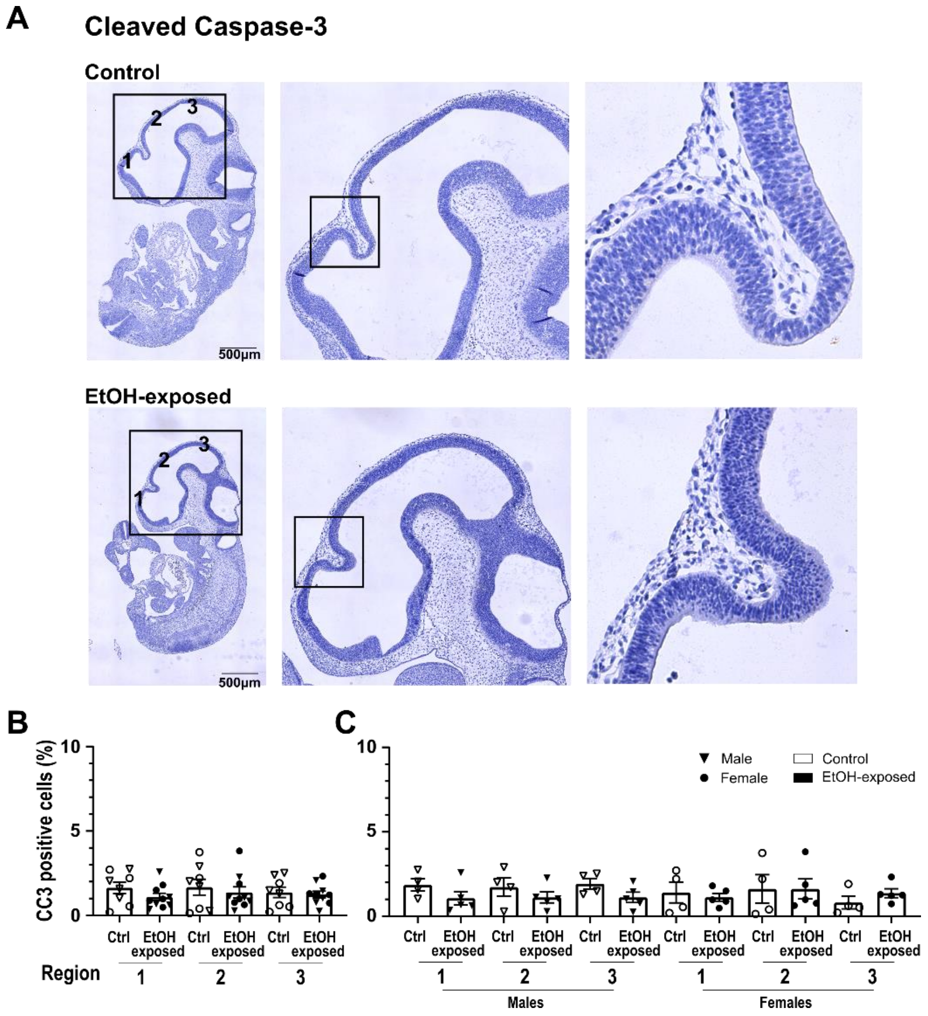


**Figure S5. Early pre-implantation alcohol exposure does not increase brain cell apoptosis in mid-gestation embryos**. Immunohistochemical staining for cleaved Caspase-3 (brown) and counterstaining with hematoxylin (blue) in control and ethanol-exposed embryos. A) Representative distribution of cleaved Caspase-3 (CC3) positive cells in sagittal sections of E10.5 embryos. Left panel shows the three brain regions used to quantify cleaved Caspase-3 positive cells. Details of framed areas are shown in middle and right panels. B) Percentage (%) of cleaved Caspase-3 positive cells across brain regions in control and ethanol-exposed (no apparent morphological defects) embryo (males and females). Bars represent mean ± SEM. C) Sex-specific quantification of cleaved Caspase-3 positive cells. Data from B) separated in male and female samples; male (Ctrl n= 4, EtOH-exposed n=5), female (Ctrl n= 4, EtOH-exposed n=5). Bars represent mean ± SEM.

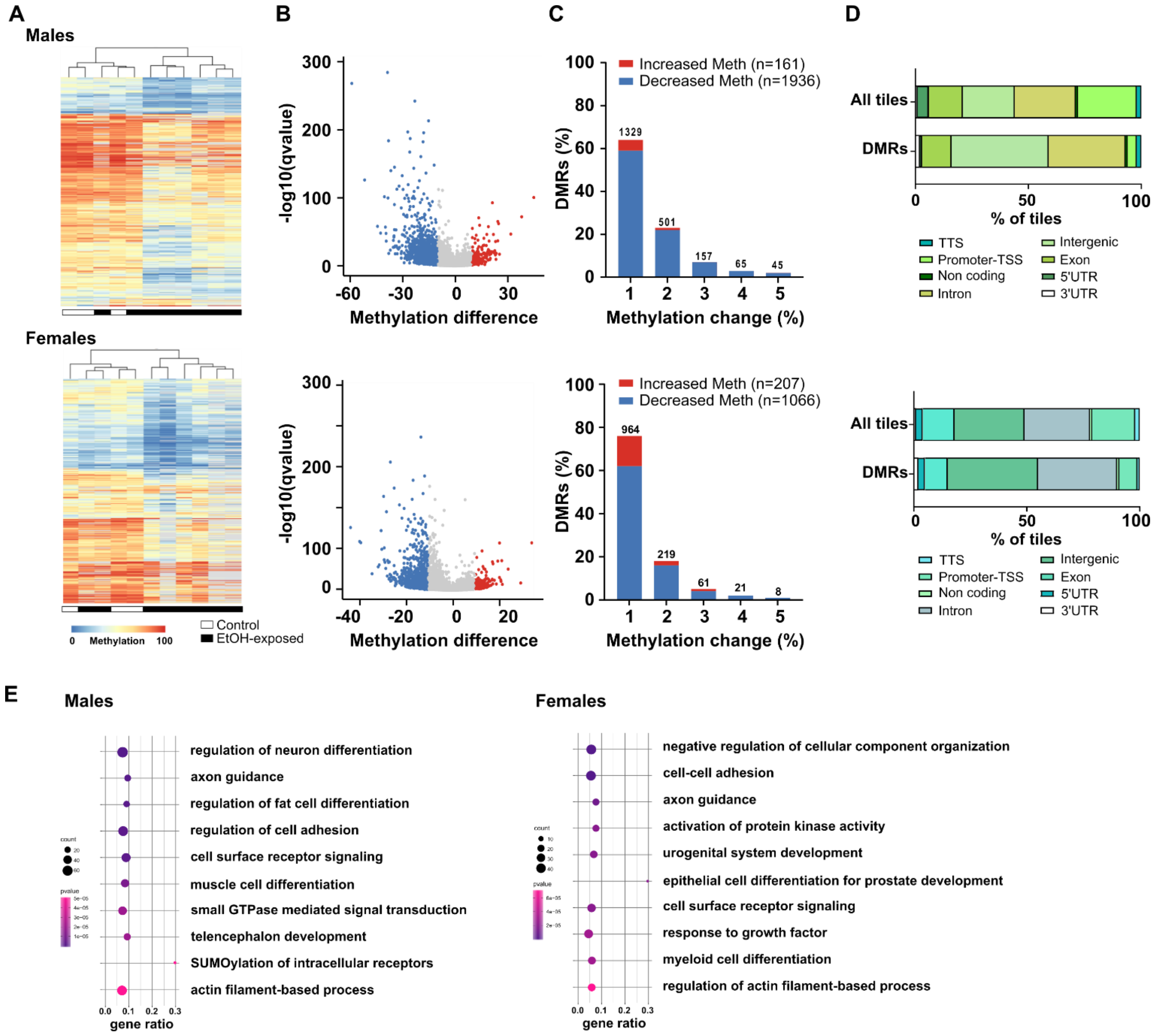


**Figure S6. Early pre-implantation alcohol exposure triggers DNA methylation alterations in developing embryonic forebrain of males and females.** Genome-wide CpG methylation analyses of male E10.5 control (n=3) and ethanol-exposed (n=8) forebrains (upper graphs) and females E10.5 control (n=3) and ethanol-exposed (n=8) forebrains (lower graphs). **A)** Heatmap showing CpG methylation levels for the DMRs (Males n=2 097; Females n=1 273) between control and EtOH-exposed forebrains. **B)** Volcano plot representing the differentially methylated regions (DMRs) between control and EtOH-exposed forebrains (see methods section for details). Red dots represent the tiles with a methylation increase of at least 10% in EtOH-exposed compared to control forebrains (Males n=161; Females n=1207); blue dots represent the tiles with a methylation decrease of at least 10% in EtOH-exposed compared to control forebrains (Males n=1 936; Females n= 1 066); grey dots represent the tiles with changes less than 10% in EtOH-exposed compared to control forebrains (Males n=81 327; Females n=125 584). **C)** Proportion of DMRs associated with the changes of CpG methylation levels between control and EtOH-exposed E10.5 forebrains. **D)** Pie chart of genomic annotations for All tiles (Males n=83 424; Females n=126 857) and DMRs (Males n=2 097 and Females (n=1 273).

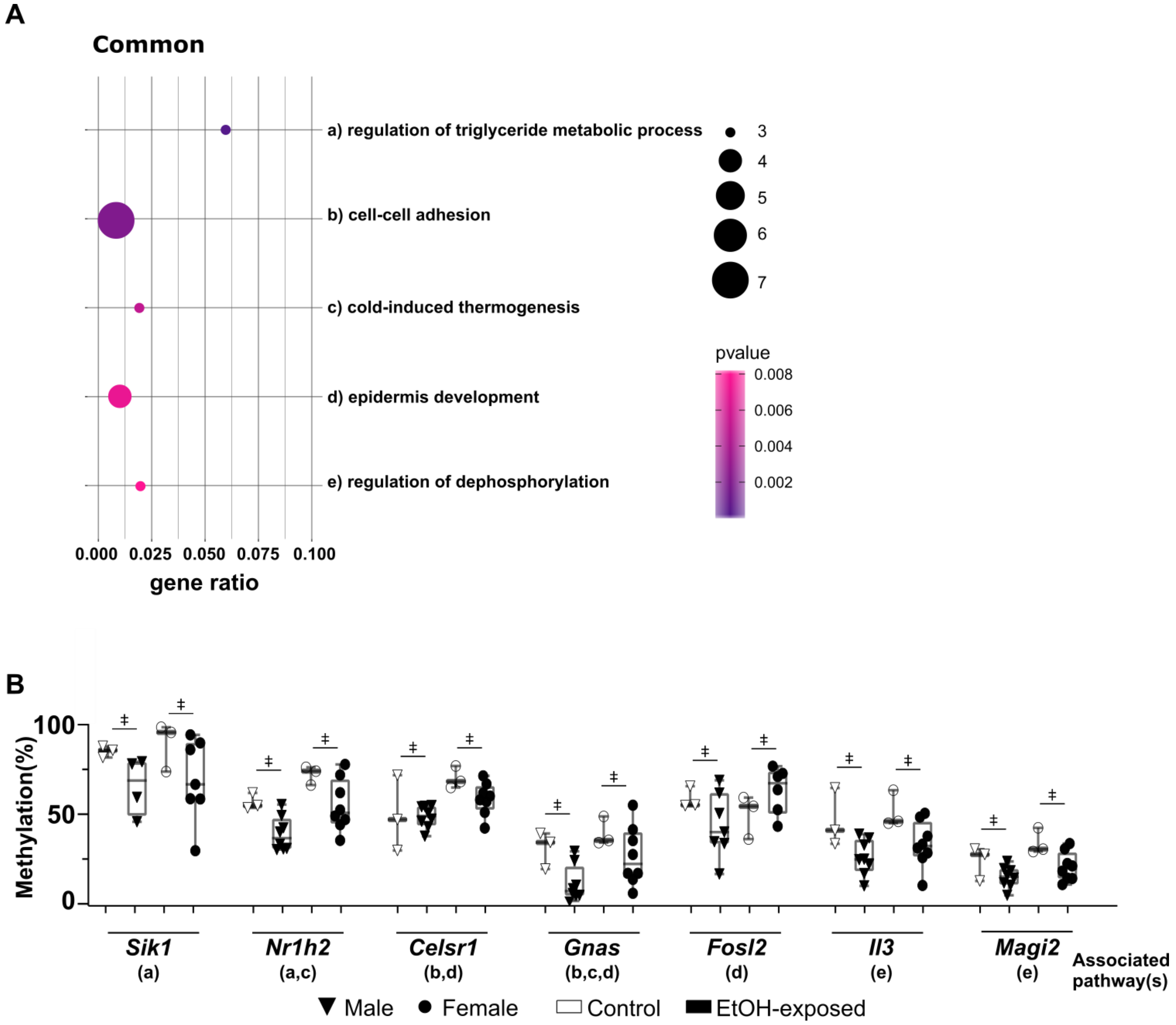


**Figure S7. Functional enrichment analysis of common DMRs. A)** Functional enrichment analysis showing the top enriched pathways for common DMRs (n=60 unique gene DMRs), based on Metascape analysis for pathways and p-value. The size of the dot represents the number of DMR-associated genes in pathways, and gene ratio represents the number of DMR-associated genes with regards to the number of genes in a pathway. **B)** Examples of CpG methylation levels of individual samples for sex-specific DMR-associated genes related to the top enriched pathways in A). Letters under gene name relate to the pathways in A). ǂ represents significant differences in CpG methylation levels of DMRs (e.g., ±>10% methylation difference, q<0.01) between control and EtOH-exposed embryos (see methods section for details).

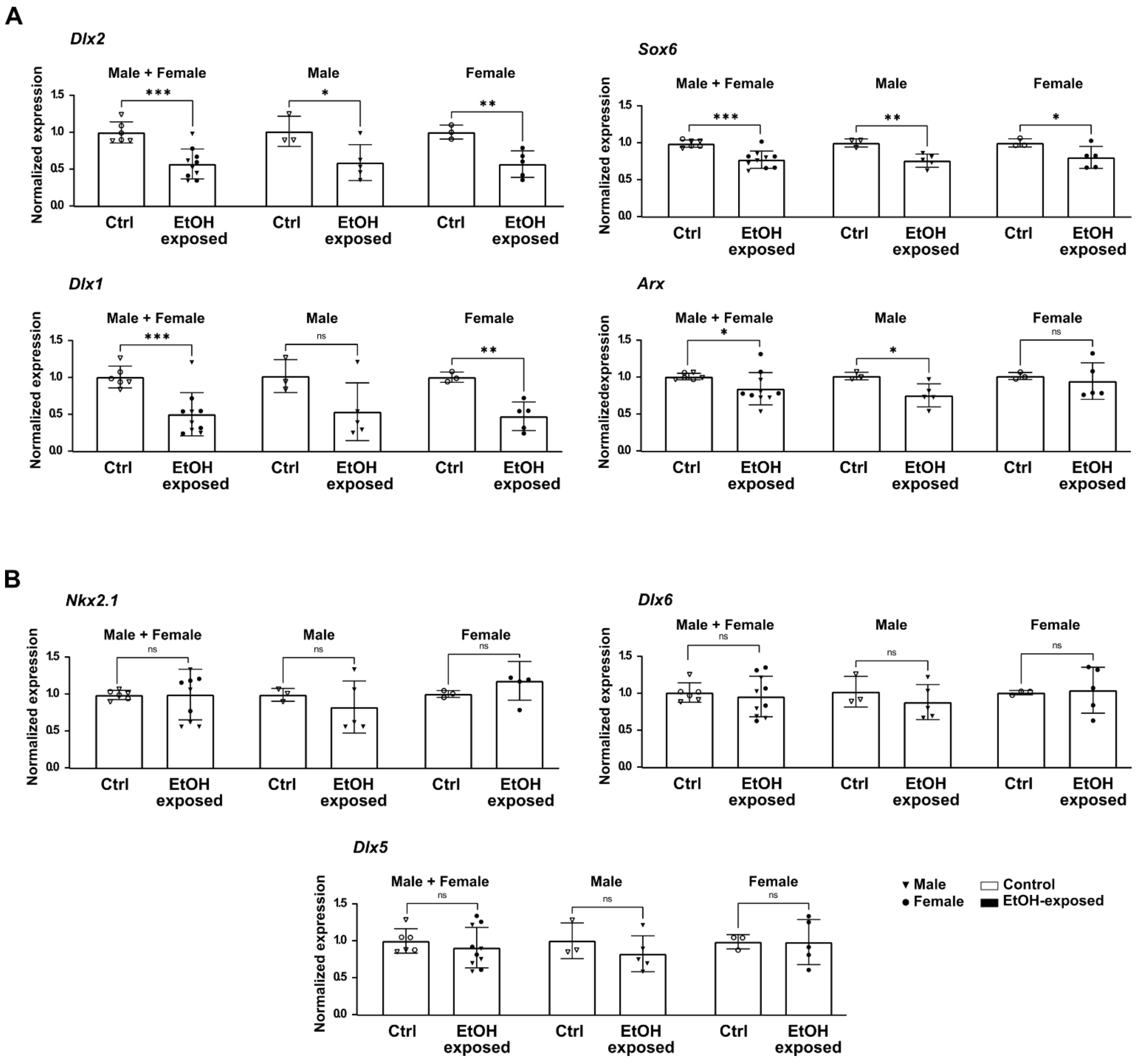


**Figure S8. Relative gene expression in forebrains following early embryonic ethanol exposure.** Selected genes associated, directly or indirectly, to the *Dlx* family of homeodomain transcription factors, which is at the core of the gene regulatory network that controls general aspects in the development of GABAergic interneurons (50,51). Quantitative Real Time PCR (qPCR) analyses of E10.5 control and EtOH-exposed forebrains showing : A) altered expression for genes associated to DMRs (*Dlx2*, *Sox6*, *Arx*) and normal DNA methylation profiles (*Dlx1*), and B) normal gene expression and normal DNA methylation profiles (*Nkx2.1*, *Dlx6*, *Dlx5*). Samples used for quantification: male Ctrl n= 3 and EtOH-exposed n=5; female Ctrl n= 3 and EtOH-exposed n=5. Gene expression was normalized using control genes *Hprt1* and *Pgk1*. Bars represent mean ± SD. * p≤0.05, ** p≤0.01, ***p≤0.001, ns: non significative.

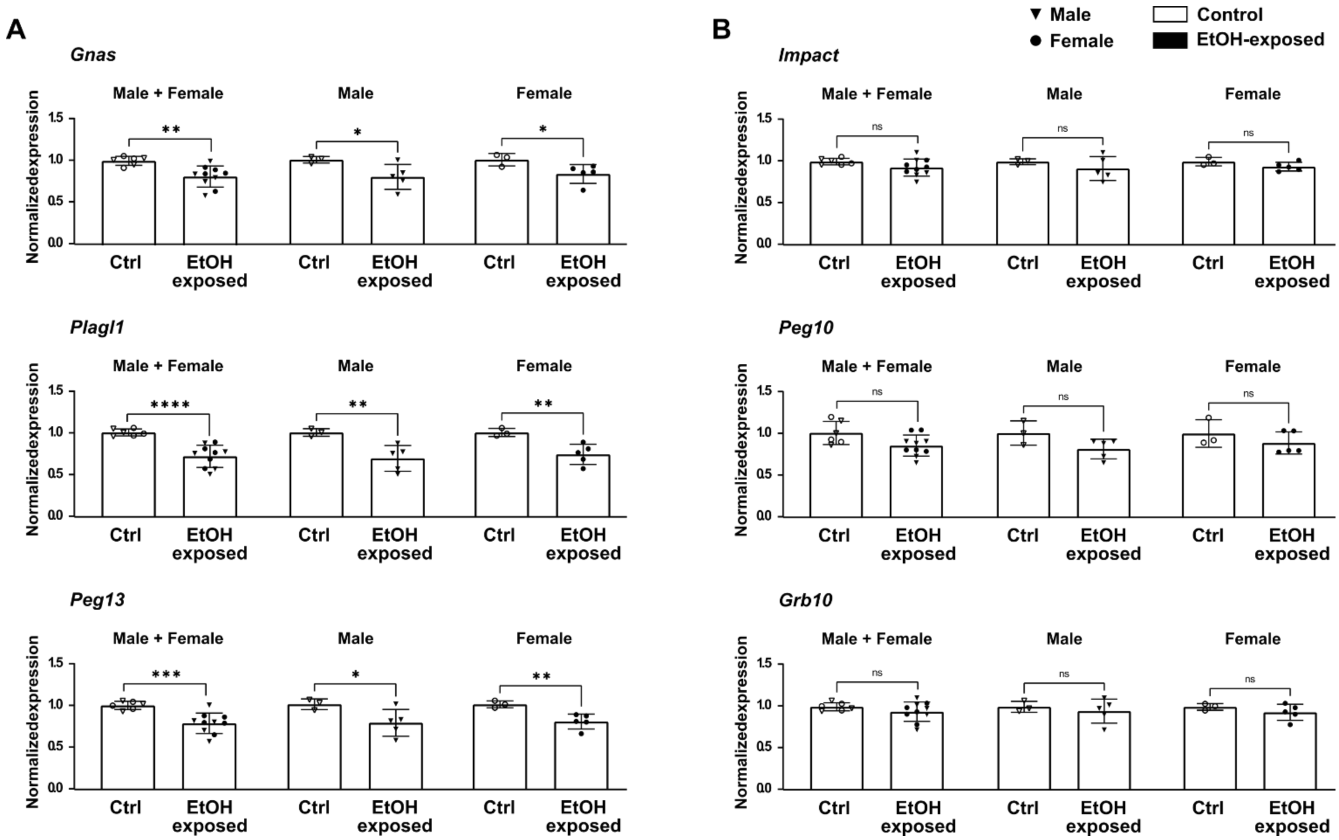


**Figure S9. Relative expression of imprinting genes in forebrains following early embryonic ethanol exposure.** Selection of imprinted genes that presented DNA methylation differences (i.e., DMRs) in their ICRs in both males and females E10.5 EtOH-exposed forebrains. Quantitative Real Time PCR (qPCR) analyses of E10.5 control and EtOH-exposed forebrains showing : A) altered expression, and B) no alteration in gene expression. Samples used for quantification: male Ctrl n= 3 and EtOH-exposed n=5; female Ctrl n= 3 and EtOH-exposed n=5. Gene expression was normalized using control genes *Hprt1* and *Pgk1*. Bars represent mean ± SD. * $p \leq 0.05$, ** $p \leq 0.01$, *** $p \leq 0.001$, **** $p \leq 0.0001$, ns: non significative.

**Table S1. Sequencing information for RRBS data.**

| Sample | #Raw reads | Alignment % | #CpGs at 10X |
|---|---|---|---|
| Ctl-1 | 21 997 747 | 75 | 974 474 |
| Ctl-2 | 20 433 987 | 78 | 1 500 815 |
| Ctl-3 | 26 320 922 | 75 | 719 769 |
| Ctl-4 | 23 030 420 | 79 | 1 066 382 |
| Ctl-5 | 24 468 312 | 79 | 965 737 |
| Ctl-6 | 27 783 216 | 79 | 1 058 976 |
| EtOH-1 | 19 896 764 | 84 | 766 792 |
| EtOH-2 | 30 101 522 | 81 | 1 066 501 |
| EtOH-3 | 24 052 738 | 80 | 932 801 |
| EtOH-4 | 26 671 945 | 84 | 859 007 |
| EtOH-5 | 24 362 110 | 80 | 1 019 589 |
| EtOH-6 | 28 346 140 | 80 | 1 061 420 |
| EtOH-7 | 21 250 969 | 81 | 724 247 |
| EtOH-8 | 39 781 961 | 87 | 1 390 455 |
| EtOH-9 | 41 482 260 | 83 | 1 624 216 |
| EtOH-10 | 22 864 202 | 81 | 853 155 |
| EtOH-11 | 39 019 936 | 81 | 1 480 644 |
| EtOH-12 | 24 787 442 | 81 | 875 794 |
| EtOH-13 | 20 102 140 | 85 | 1 017 251 |
| EtOH-14 | 25 365 210 | 81 | 936 405 |
| EtOH-15 | 26 574 342 | 77 | 1 635 345 |
| EtOH-16 | 23 808 905 | 83 | 842 291 |

**Table S2. Primer sequences**

| Gene | Forward | Reverse |
|---|---|---|
| *Arx* | GCTCTCCTCCTACTGCATCG | CTCCCAGAAGCCTCATTTTG |
| *Dlx1* | TGTCTCCTTCTCCCATGTCC | TGCTGACCGAGTTGACGTAG |
| *Dlx2* | TGGGCTCCTACCAGTACCAC | TTGTTGACCGGAGACGAACT |
| *Dlx5* | TCTCAGGAATCGCCAACTTT | GAGCGCTTTGCCATAAGAAG |
| *Dlx6* | GGAGGCAACTCCTACAACCA | GACTGGAGGTAAGGGCTGTG |
| *Gnas* | CATTCTGAGCGTGATGAACG | ATCCTCCCACAGAGCCTTG |
| *Grb10* | GCTTCTCCATCTGTGAAGTGG | AAGACCACTGCGAGATTTTCA |
| *Hprt1* | CTGGTGAAAAGGACCTCTCGAA | CTGAAGTACTCATTATAGTCAAGGGCAT |
| *Impact* | ACTTCATGGATGACCCCAAAT | TGATAAGAAGGCGGTGCTGTA |
| *Nkx2.1* | AAAGCACACGACTCCGTTCT | CTCCATGCCCACTTTCTTGT |
| *Peg10* | GGACCCCTCATCCTTCGT | GTTGGCGTCTTTTGGTTCTT |
| *Peg13* | TAAAGTGCCCTGATCCGAAC | ATTTCAAACCTGCCAACCTG |
| *Pgk1* | CTGACTTTGGACAAGCTGGACG | GCAGCCTTGATCCTTTGGTTG |
| *Plagl1* | AATGTGGCAAGTCCTTCGTCAC | TGGTTCTTCAGGTGGTCCTTCC |
| *Sox6* | CAGCCCATGATGAACAGAAA | AGTTGCTGCTGTTGTCTTGC |
